# Transcriptomic Characterization Reveals Disrupted Medium Spiny Neuron Trajectories in Huntington’s Disease and Possible Therapeutic Avenues

**DOI:** 10.1101/2023.04.30.538872

**Authors:** Carlos Galicia Aguirre, Kizito-Tshitoko Tshilenge, Elena Battistoni, Alejandro Lopez-Ramirez, Swati Naphade, Kevin Perez, Sicheng Song, Sean D. Mooney, Simon Melov, Michelle E. Ehrlich, Lisa M. Ellerby

## Abstract

Huntington’s disease (HD) is a neurodegenerative disorder caused by an expansion of CAG repeats in exon 1 of the *HTT* gene, ultimately resulting in the generation of a mutant HTT (mHTT) protein. Although mHTT is expressed in various tissues, it significantly affects medium spiny neurons (MSNs) in the striatum, resulting in their loss and the subsequent motor function impairment in HD. While HD symptoms typically emerge in midlife, disrupted MSN neurodevelopment has an important role. To explore the effects of mHTT on MSN development, we differentiated HD induced pluripotent stem cells (iPSC) and isogenic controls into neuronal stem cells, and then generated a developing MSN population encompassing early, intermediate progenitors, and mature MSNs. Single-cell RNA sequencing revealed that the developmental trajectory of MSNs in our model closely emulated the trajectory of fetal striatal neurons. However, in the HD MSN cultures, the differentiation process downregulated several crucial genes required for proper MSN maturation, including Achaete-scute homolog 1 and members of the DLX family of transcription factors. Our analysis also uncovered a progressive dysregulation of multiple HD-related pathways as the MSNs matured, including the NRF2-mediated oxidative stress response and mitogen-activated protein kinase signaling. Using the transcriptional profile of developing HD MSNs, we searched the L1000 dataset for small molecules that induce the opposite gene expression pattern. Our analysis pinpointed numerous small molecules with known benefits in HD models, as well as previously untested novel molecules. A top novel candidate, Cerulenin, partially restored the DARPP-32 levels and electrical activity in HD MSNs, and also modulated genes involved in multiple HD-related pathways.

## INTRODUCTION

Huntington’s disease (HD) is a fatal, dominantly inherited neurodegenerative disorder that primarily affects neurons in the striatum and cortex [1, 2]. HD is caused by a CAG expansion in the huntingtin gene that leads to a polyglutamine (polyQ) expansion in the encoded protein (HTT), and patients with a CAG expansion greater than 38 repeats exhibit symptoms [3]. As a basal ganglia disease, HD impacts motor learning and control (chorea), executive functions, and emotions. The major cell type and principal output neuron of the caudate-putamen is the inhibitory g-aminobutyric acid (GABA)-ergic medium spiny neurons (MSNs). Dysfunction and developmental alterations of MSNs have been implicated in HD, with MSN subtypes being differently impacted [4-14]. Despite the mutant HTT (mHTT) protein being present throughout life, HD symptoms only manifest later in life. This raises the question of whether the early developmental alterations in MSNs set the stage for developing HD symptoms later in life. As seen in human imaging studies, pre-onset young HD carriers already have abnormal striatal volumes [15, 16]. These observations have been know for a number of years, [17] and arose from findings of abnormalities in immature cells carrying the mutation [18, 19]. Humbert and colleagues [4] showed clear abnormalities in cortical development in human fetuses with mHTT and in engineered mice, with many deficits in cell cycling, neuronal differentiation, and endosome dynamics. The KIDS-HD study (reviewed in [20]), including studies that focused on the developmental trajectory of the striatum, demonstrated hypertrophy in the early years of childhood and then a steep decline. These results are in line with studies conducted on HD developmental models, that showed an increase in neuronal proliferation and premature maturation [21, 22].

We reported isogenic HD neuronal stem cells (NSCs) derived from induced pluripotent stem cells (iPSCs) display dysregulated signaling pathways [6, 23, 24]. Specifically, we used human patient–derived HD-iPSCs (72CAG/19CAG, HD72) and genetically corrected the cells to a normal repeat length (21CAG/19CAG, C116), thus creating an isogenic control [23]. Our transcriptomic analysis of isogenic HD72 NSCs suggested that HD is linked to developmental impairments that prevent proper generation of MSNs and subsequent loss of MSNs identity [6, 8, 23-25]. Similarly, HD iPSC derived neurons feature dysregulation of multiple pathways related to development [21, 26, 27]. We also found that deletion of transcription factors required for striatum development leads to HD-like phenotypes [28]. This highlights the importance of correct striatum development and suggests that targeting this pathway has important therapeutic implications for HD.

In studies in mice, transient expression of mHTT during development leads to neurodegeneration and HD-like symptoms later in life [7]. Furthermore, lower levels of the non-mutant HTT during development result in HD-like motoric abnormalities in adulthood [29]. Conversely, correcting early neuronal defects caused by mHTT in mice can delay the onset of HD pathology [12]. These findings highlight the potential for treating developmental deficits in HD as a therapeutic approach and underscore the importance of understanding the mechanisms of dysregulation caused by the mutation during early stages of development.

Since the HD gene, *HTT*, was discovered in 1993 [30], progress has been made in understanding the cellular pathways disrupted in HD and in identifying potential therapeutics. Still, the proximal events in HD pathogenesis remain unclear and there are no treatments modify the disease progression. Thus, a comprehensive characterization of the very early proximal events in HD MSNs is critical for the design of alternative therapeutic approaches to delay disease onset and progression.

In this study, we investigated the effects of mHTT on the developmental trajectory of MSNs. We utilized our isogenic HD72 and C116 iPSC lines [23] to generate developing MSNs at various stages of maturation, including early and intermediate progenitors and mature MSNs. To comprehensively analyze the developmental changes, we used bulk and scRNA sequencing. By integrating our data with scRNAseq data of the developing human striatum [31], we identified the developmental trajectories affected by HD. Our analysis revealed that multiple pathways were disrupted during HD72 MSN development, along with alterations in transcription factors responsible for MSNs identity. Notably, Achaete-scute homolog 1 (*ASCL1*) and the Distal-less homeobox (*DLX*) family of transcription factors, essential for MSN development, were downregulated in developing HD72 MSNs and we also see downregulation in adult HD patients and post-natal mouse models. With the transcriptomic data, we identified potential therapeutics, including several small molecules known to improve HD phenotypes, and untested molecules. We assessed the effects of Cerulenin, a small molecule not previously used in HD, and found a partial rescue of DARPP-32 levels and, electrical activity and modulation of multiple HD-related pathways in HD72 MSNs.

## RESULTS

### Differentiation and Characterization of HD72 NSCs and Developing MSNs

We differentiated HD72 and C116 iPSCs into NSCs using a monolayer differentiation approach [32]. (**Fig. 1A**). After confirming the expression of the NSC marker nestin (NES) (**Fig. 1B**) [33], the cells were differentiated into a mixed population of early, and intermediate progenitors, and mature MSNs using a chemical protocol that mimics the developmental stages of the lateral ganglionic eminence (LGE) [34]. After 21 days of differentiation, DARPP-32 positive cells were visible among the developing MSNs (**Fig. 1C**). Furthermore, developing MSNs expressed higher levels of dopamine and cAMP-regulated phosphoprotein Mr 32,000 (*DARPP-32, PPP1R1B*), calbindin 1 (*CALB1*), calbindin 2 (*CALB2*), dopamine receptor D1 (*DRD1*), dopamine receptor D2 (*DRD2*), BCL11 transcription factor B (*CTIP2*), opioid receptor mu 1 (*OPRM1*, also known as *MOR1-6TM*, *MOR1-7TM*), and nuclear receptor subfamily 4 group A member 1 (*NR4A1*) mRNA compared to NSCs (**Fig. 1D**).These results indicate the presence of mature MSNs in the population.

**Figure 1.**
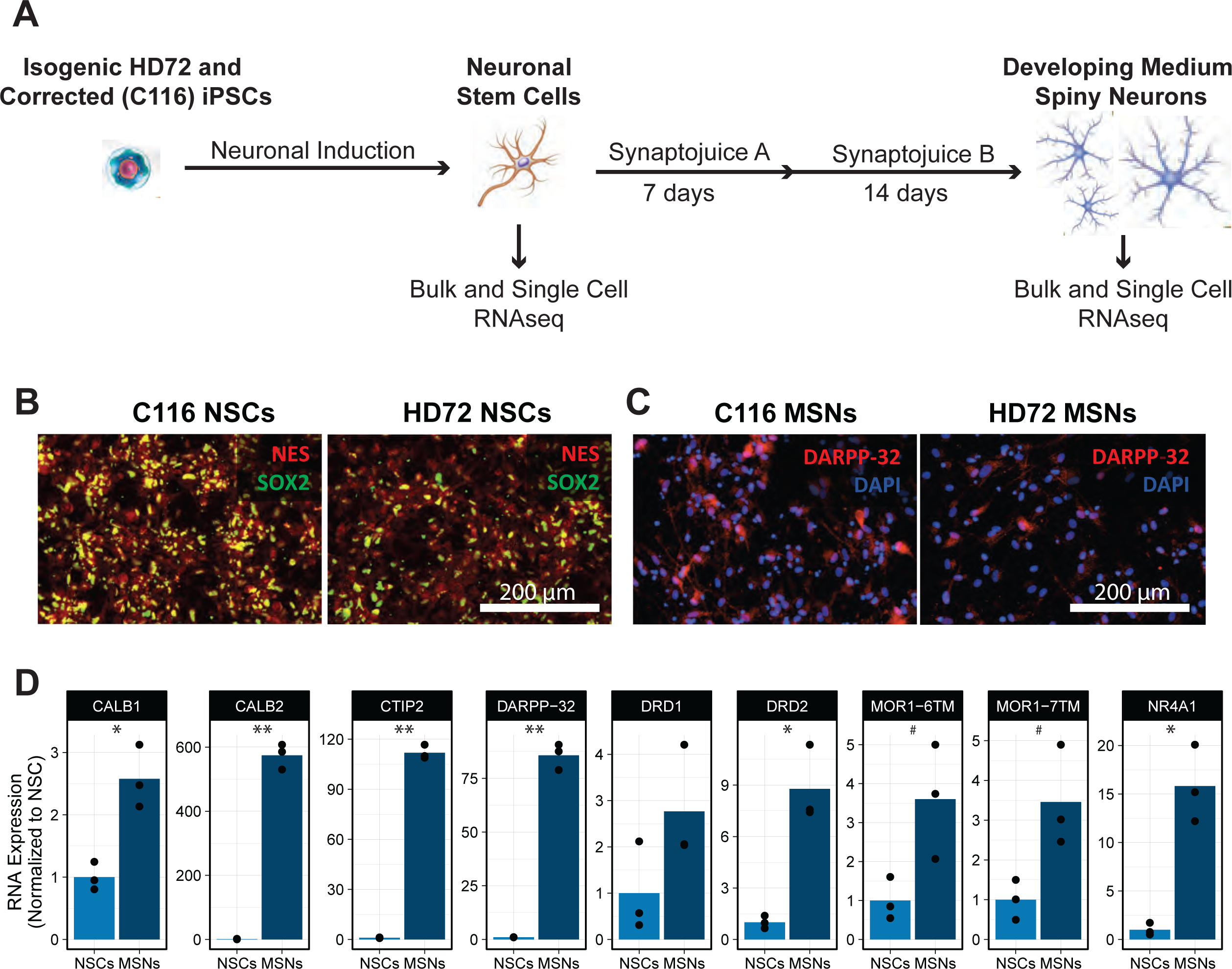
Differentiation of Developing MSNs. **(A)** Diagram of differentiation process for MSNs. **(B)** C116 and HD72 NSCs immunolabeled with Nestin and SOX2**. (C)** DARPP-32 staining on developing MSNs. Scale bar: 200 μm. **(D)** RNA expression of multiple markers of MSNs in NSCs and developing MSNs determined with qPCR. P values were calculated using t tests followed by Benjamini-Hochberg correction for multiple tests. # < 0.1, * < 0.05, ** < 0.01.

Using bulk RNA sequencing we compared the transcriptional profiles of HD72 developing MSNs to a published RNAseq dataset of HD72 NSCs [6]. Principal component analysis (PCA) showed a distinct separation between HD72 and C116 developing MSNs (**Fig. 2B**), whereas HD72 and C116 NSCs clustered more closely (**Fig. 2A**). We depicted the transcriptional changes induced by mHTT in NSCs and developing MSNs using volcano plots (**Fig. 2C, D**). A total of 4,533 and 4,082 genes were differentially expressed in MSNs and NSCs, respectively (adjusted p-value < 0.01, absolute log2-fold change > 0.1) (**Supplemental Table 1 and 2**). Notably, in HD72 NSCs, the top downregulated differentially expressed genes (DEGs) included *MSH* homeobox 1 (*MSX1*), a modifier of age of onset in HD [35]. In HD MSNs, we observed a notable upregulation of insulin-like growth factor binding protein 7 (*IGFBP7*), a gene implicated in the initiation of senescence and apoptosis [36]. These findings align with our previous discovery that HD MSNs display numerous senescence-like characteristics [37]. Though MSNs and NSCs shared 1,362 DEGs, their correlation was low (Pearson’s r = 0.06, p-value < 2.2e-16) (**Supplemental Fig. 1A**). As anticipated, since MSNs are the main cell type affected by HD, DEGs in MSNs demonstrated a larger fold change compared to those in NSCs.

**Figure 2.**
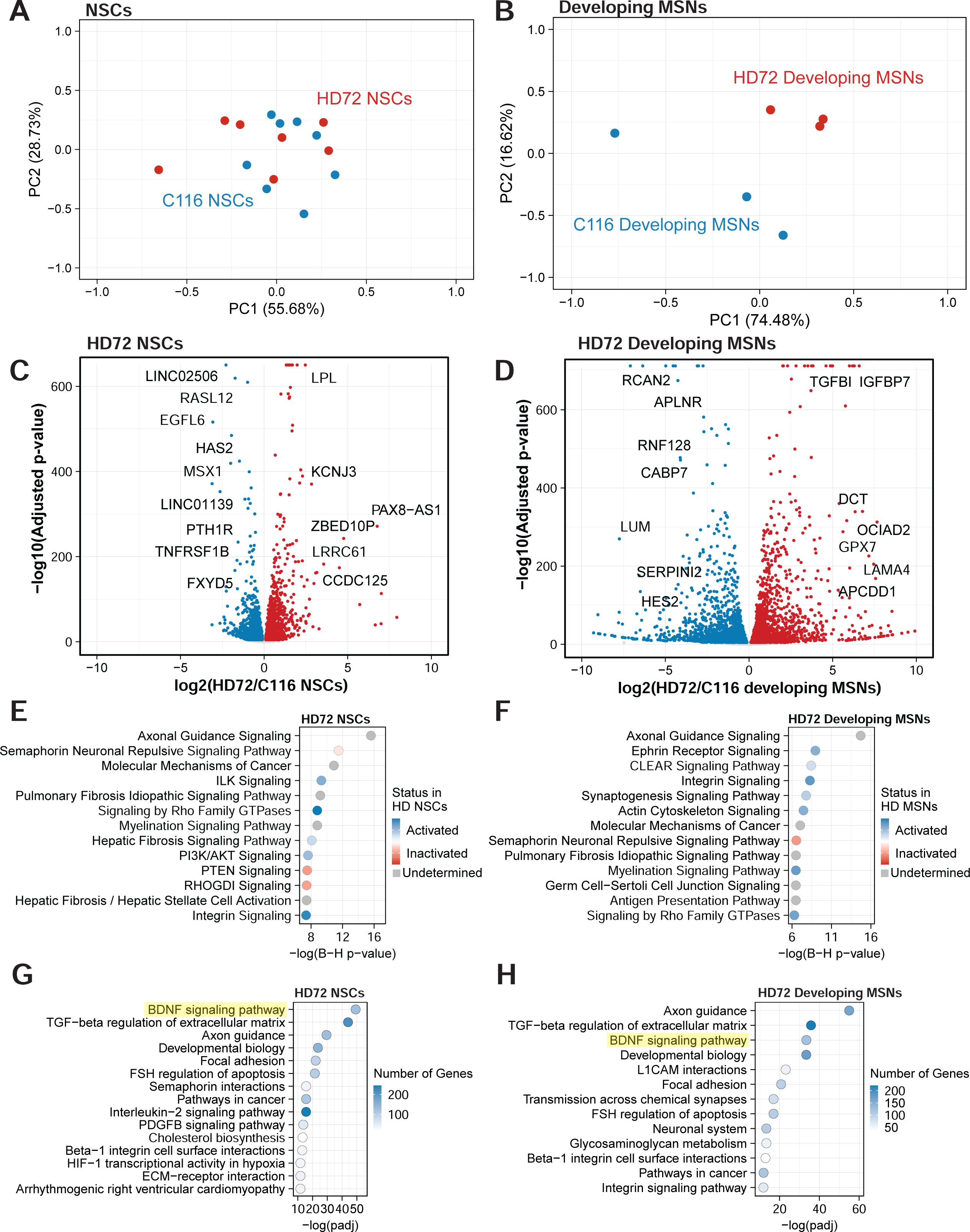
Transcriptional characterization of HD72 NSCs and developing MSNs. **(A,B)** PCA plot of NSCs (A) and developing MSNs (B). **(C,D)** Volcano plots illustrating DEGs when comparing HD72 vs C116 NSCs (C) and HD72 MSNs vs C116 MSNs (D). **(E,F)** IPA of HD72 NSCs (E) and HD72 MSNs (F). Gray dots indicate an undetermined direction for the pathway. **(G,H)** Bioplanet enrichment analysis of HD72 NSCs (G) and HD72 MSNs (H).

Next, we sought to assess whether RNA alterations were reflected at the protein level in developing HD72 MSNs. We compared our RNAseq data with our recently published proteomics dataset from developing HD72 and C116 MSNs [38]. We identified a considerable correlation between gene fold change values at both the RNA and protein levels (Pearson’s r = 0.75, p-value < 2.2e-16), highlighting the potential impact of these alterations on cellular function (**Supplemental Fig. 1B**). We also compared the DEGs in developing HD72 MSNs to those found in other HD models and data from HD patients. Our comparisons encompassed the striatum of BACHD-ΔN17 at 2, 7, and 11 months of age [39]; the striatum of Q80, Q92, Q111, Q140, and Q175 at 2, 6, and 10 months [40]; the striatum from 11-week-old R6/2 mice [41]; DEGs from iPSC-derived MSNs [42]; and data from the caudate and putamen of HD patients [43]. We noted a significant overlap in DEGs across most comparisons (**Supplemental Table 2**). As anticipated, the highest similarity was observed with iPSC-derived MSNs (adjusted p-value 1.6E-35). The mouse model exhibiting the greatest similarity was the 11-week-old R6/2 (adjusted p-value 2.16E-220). In adult HD mice this likely represents the dedifferentiation of MSNs [44]. Additionally, a significant overlap was detected with data obtained from Grade 2 HD caudate postmortem tissue (adjusted p-value 1.70E-50).

We employed Ingenuity Pathway Analysis (IPA) to assess alterations in canonical pathways within HD72 NSCs and developing MSNs (**Supplemental Table 1 and 2**). The top IPA predictions in HD72 NSCs included, dysregulation of axonal guidance and molecular mechanisms of cancer, along with activation of integrin-linked kinase (ILK) signaling, and ras homolog (RHO) family GTPase signaling. Additionally, we observed inactivation of semaphorin-mediated neuronal repulsion, phosphatase and tensin homolog (PTEN) signaling, and rho GDP dissociation inhibitor alpha (RHOGDI) signaling (**Fig. 2E**). In developing MSNs, top IPA predictions included dysregulation of axonal guidance and molecular mechanisms of cancer, activation of ephrin receptor, integrin, and coordinated lysosomal expression and regulation (CLEAR) signaling pathways, along with inactivation of semaphorin-mediated neuronal repulsion, RHOGDI signaling, and the sumoylation pathway (**Fig. 2F**). Furthermore, enrichment analysis using NCATS BioPlanet identified multiple HD-associated pathways (**Supplemental Table 1 and 2**). Among the top terms enriched in HD72 NSCs and developing HD72 MSNs were axon guidance, transforming growth factor (TGF-β) regulation of extracellular matrix, developmental biology, brain derived neurotrophic factor (BDNF) signaling and focal adhesion [6, 45-47] (**Fig. 2G, H**). Distinct enrichments in developing HD72 MSNs, which were not present in HD72 NSCs, included transmission across chemical synapses, neuronal systems, and lysosomes, among others (**Supplemental Table 2**). Importantly, the BDNF pathway has been found to be dysregulated in the striatum of HD patients, various cellular models expressing mHTT, and multiple mouse models including Hdh ^109/109^ kock-in, YAC, N-171 82Q, and R6/2 among others [48].

We used Leafcutter [49] to identify 248 genes that had differential RNA-splicing in developing HD72 MSNs (**Supplemental Table 3**). We compared these genes with those mis-spliced in the striatum of human HD patients and the R6/1 HD model [50], revealing 133 and 40 shared mis-spliced genes, respectively (**Supplemental Fig. 1C**). Enrichment analysis of the mis-spliced genes common to all three datasets demonstrated an overrepresentation of axon guidance and long-term potentiation pathways. Notably, both pathways are dysregulated in HD [46, 51] (**Supplemental Fig. 1D**, **Supplemental Table 3**).

### scRNAseq Reveals Dysregulation of Pathways Related to HD Pathology as MSN Maturation Occurs

We performed scRNAseq on developing HD72 and C116 MSNs to explore transcriptional dysregulation at single-cell resolution during HD MSN development. Our analysis revealed a continuous maturation trajectory encompassing i) early progenitors, identified by the expression of *NES*, and Vimentin (*VIM*); ii) intermediate progenitors, characterized by ASCL1, Sp9 transcription factor (*SP9*), *DLX1* and *DLX2* expression; and iii) mature MSNs, distinguished by forkhead box P1 (FOXP1), early B cell factor-1 (*EBF1*), and microtubule associated protein 2 (MAP2) (**Fig. 3A, B**). All these clusters expressed *MEIS2*, indicating an LGE-like lineage. Notably, despite DARPP-32 being detected at the protein level and RNA level via qPCR and bulk RNAseq, it was not well detected in scRNAseq, presumably due to low abundance leading to increased dropout. Similar constrains have previously been observed in scRNA studies of the developing striatum [31].

**Figure 3.**
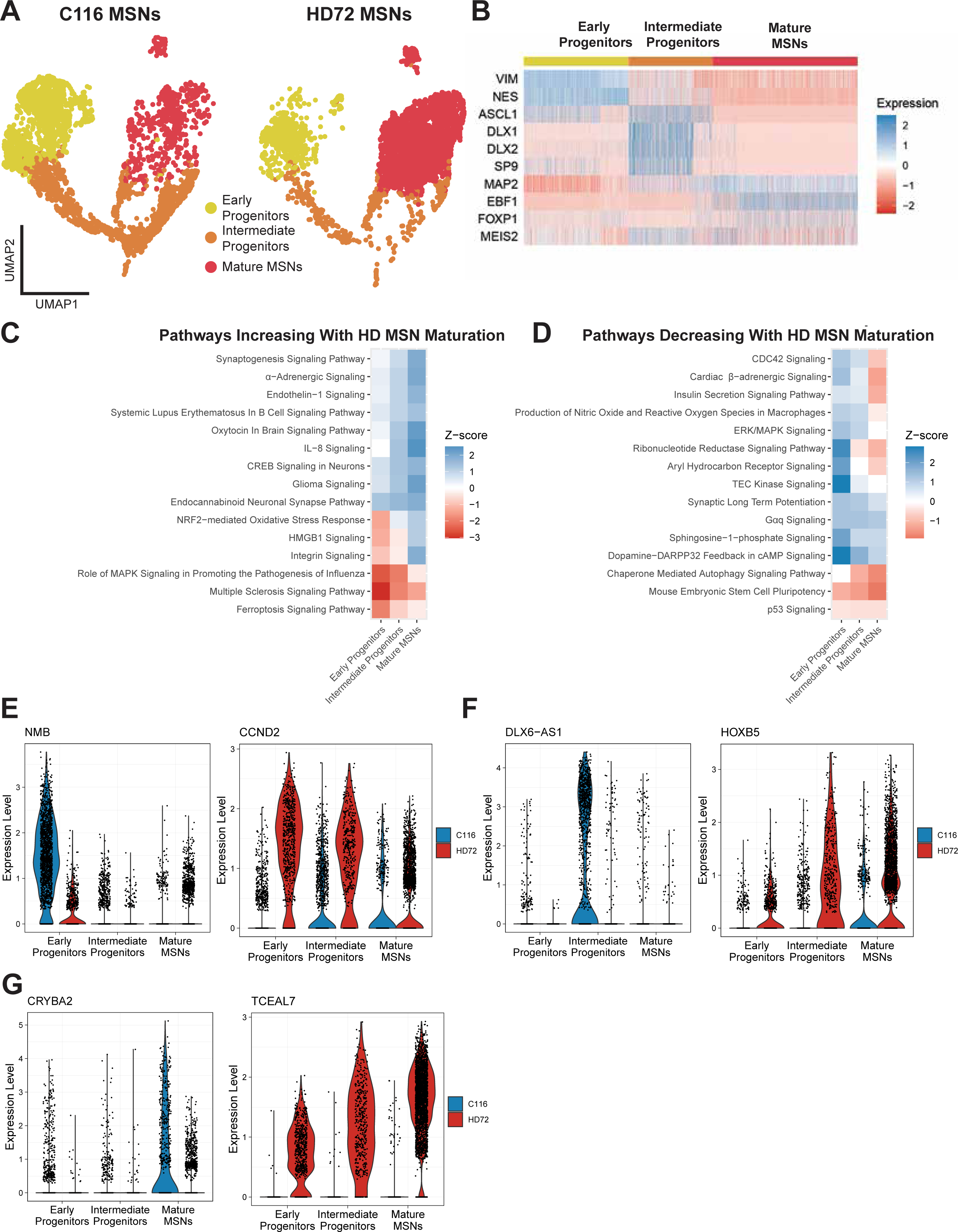
Single Cell Transcriptional Characterization of HD72 and C116 Developing MSNs. **(A)** UMAP plots of C116 and HD72 MSNs. **(B)** Heatmap showing levels of expression of various markers of early progenitors, intermediate progenitors and mature MSNs. **(C)** Heatmap showing level of activation of pathways that become more activated with MSNs maturation. **(D)** Heatmap showing level of activation of pathways that become less activated with MSNs maturation. **(E)** Violin plots showing expression of the top upregulated and downregulated genes in HD72 early progenitors, NMB and CCND2. **(F)** Violin plots showing expression of the top upregulated and downregulated genes in HD72 intermediate progenitors, GPC1 and DHRS3. **(G)** Violin plots showing expression of the top upregulated and downregulated genes in HD72 intermediate progenitors, DLX6-AS1 and TCEAL7.

Next, we conducted differential gene expression analysis between C116 and HD72 cells at each developmental stage and utilized IPA analysis to predict the activation state of canonical pathways (**Supplemental Table 4**). Our results revealed that, as MSNs matured, several pathways associated with HD pathology exhibited increased activation (**Fig. 3C**). Specifically, α-adrenergic signaling was predicted to increase, which is consistent with prior research linking α-adrenergic receptors to heightened neurotoxicity in HD [52]. Additionally, the nuclear factor erythroid 2-related factor 2 (NRF2)-mediated oxidative stress response, which is increased in HD in response to oxidative stress, displayed increased activation with MSN maturation [53]. In contrast, we also observed decreased activation in pathways that are protective against HD as HD72 MSNs matured (**Fig. 3D**). For example, activation of both sphingosine-1-phosphate and mitogen-activated protein kinase 1 (ERK/MAPK) signaling diminished with increased maturation, and their activation has been associated with exerting protective effects in HD [54, 55]. Additionally, we observed a reduction in Dopamine-DARPP32 feedback within cAMP signaling, a pathway known to be downregulated in HD [56]. Among the top differentially expressed genes, we discovered neuromedin B (*NMB*) and cyclin D2 (*CCND2*) in early progenitors (**Fig. 3E**). In intermediate progenitors, the most dysregulated genes included DLX6 antisense RNA 1 (*DLX6-AS1*) and homeobox B5 (*HOXB5*) (**Fig. 3F**). Meanwhile, in mature MSNs, crystallin beta A2 (*CRYBA2*) and transcription elongation factor A-like 7 (*TCEAL7*) emerged as the top differentially expressed genes (**Fig. 3G**). Notably *DLX6-AS1* interacts with other DLX transcription factors important during striatum development [57].

We also used scRNAseq in C116 and HD72 NSCs to show heterogeneous expression of multiple genes. We separated them into four clusters and displayed the top genes representing their heterogeneity (**Supplemental Fig. 2A, B**). Despite their heterogeneity, they showed homogeneous expression of NSC marker NES indicating a single cell type (**Supplemental Fig. 2C**). Differential expression in C116 and HD72 NSCs showed among the top downregulated genes in HD72 NSCs, Ubiquitin B (*UBB*), Retinol binding protein 1 (*RBP1*) and Crystallin beta B1 (*CRYBB1*). Among the top upregulated genes in HD72 NSCs, we found OCIA domain containing 2 (*OCIAD2*), insulin like growth factor binding protein 5 (*IGFBP5*) and S100 calcium binding protein B (*S100B*) (**Supplemental Fig. 2D, Supplemental Table 4**). Interestingly, OCIAD2 is linked to Alzheimer’s [58], and IGFBP5 induces cell senescence [59]. Previously, we’ve observed senescence-like traits in HD72 MSNs [37].

### HD Alters the Neurodevelopmental Program of MSNs

To compare the in vitro maturation of MSNs with their native human development, we analyzed scRNAseq data from our model after integrating it with a human LGE dataset spanning weeks 7, 9, and 11 post-conception (**Fig. 4A,B**) [29]. During fetal development, MSNs begin as apical progenitors, progress to basal progenitors, and ultimately mature into fully differentiated MSNs. Our goal was to assess whether in vitro maturation of MSNs follows a similar trajectory by comparing the expression patterns of various markers throughout these developmental stages (**Fig. 4A-C**). We observed that *NES*, an apical progenitor marker, was expressed in both apical and early progenitors. In contrast, GS homeobox 2 (*GSX2*) and transcription factor 7 like 1 (*TCF7L1*), both markers of apical progenitors, were exclusively expressed in apical progenitors rather than in early progenitors. This finding highlights a distinct identity between these two cell populations at this particular developmental stage. On the other hand, *ASCL1* and *DLX1*, markers for basal progenitors, were present in both basal and intermediate progenitors (**Fig. 4C**). Additionally, *FOXP1*, a marker of mature MSNs, was detected in both the LGE and in vitro datasets (**Fig. 4C**). We also observed similar patterns of expressions in developing MSNs and LGE for *MEIS2* a marker of the LGE lineage, and doublecortin (*DCX*) a marker of neurons.

**Figure 4.**
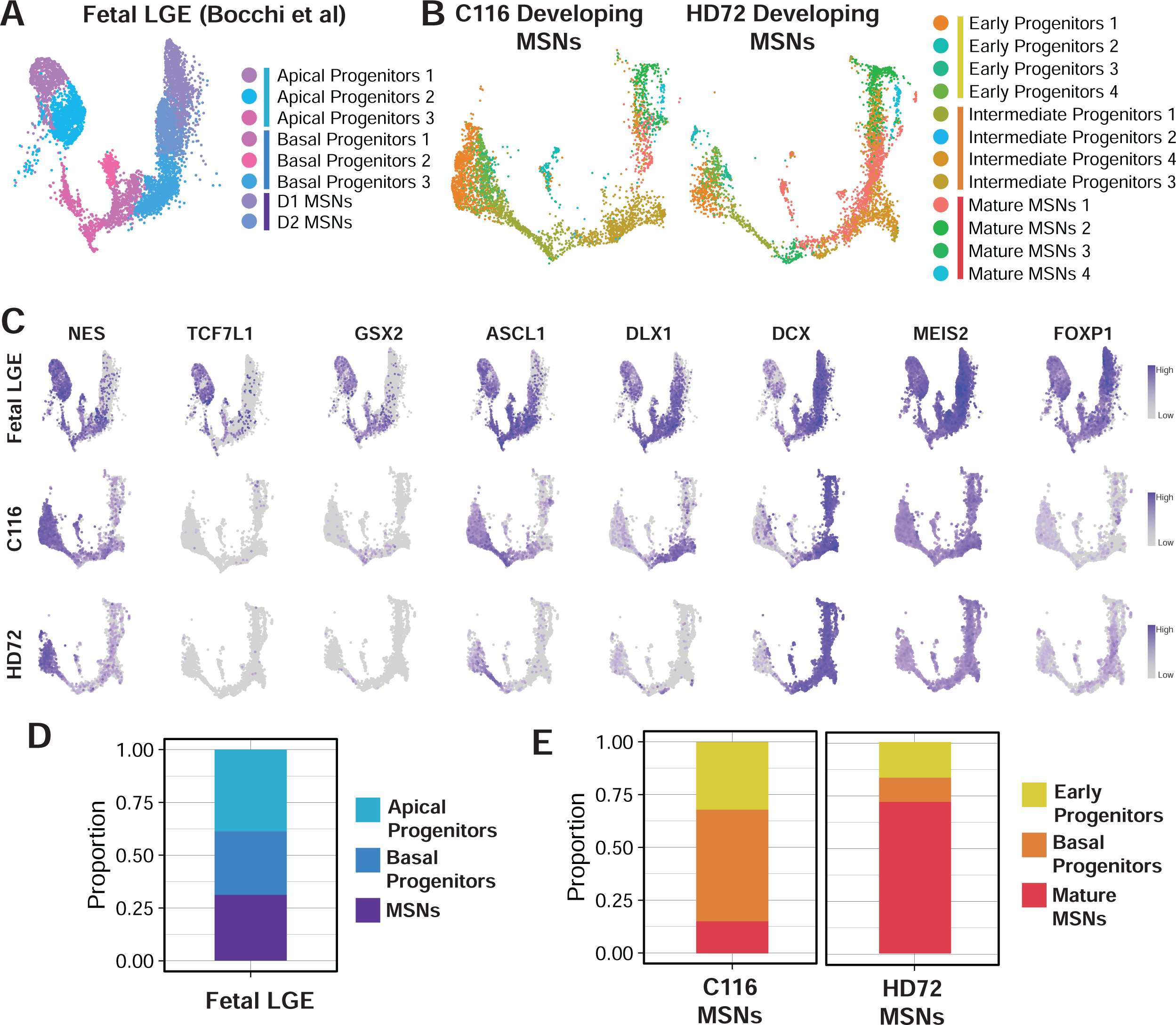
Comparison of iPSC Derived C116 and HD72 MSNs with Human Fetal LGE. **(A,B)** UMAP plot of fetal LGE (A)C116 and HD72 developing MSNs (B). **(C)** Expression of NES, TCF7L1, GSX2, ASCL1, DLX1, DCX, MEIS2 and FOXP1. **(D)** Proportion of apical progenitors, basal progenitors and MSNs in LGE. **(E)** Proportion of early progenitors, basal progenitors and mature MSNs in C116 and HD72 MSNs.

We analyzed the frequency of apical progenitors, basal progenitors, and mature MSNs in the LGE dataset, finding similar ratios for these three cell types (**Fig. 4D**). In a comparison of C116 and developing HD72 MSNs, we observed a higher percentage of mature MSNs and a lower percentage of early and intermediate progenitors in developing HD72 MSNs (**Fig. 4E**). This observation was supported by NES immunostaining, which showed reduced NES expression in developing HD72 MSNs (**Supplemental Fig 3A,B**).

Next, we aimed to characterize gene expression trajectories during MSN maturation in the three datasets. We employed VIA to model MSN maturation progression [60]. VIA builds a k-nearest neighbor graph in which nodes represent cell clusters linked according to their gene expression similarity. The algorithm takes a user-provided root node as the trajectory’s starting point and infers the trajectory using lazy-teleporting random walks, combined with Markov chain Monte Carlo refinement. To initialize the algorithm, we designated the clusters for apical and early progenitors with the highest NES expression as root nodes that, represented the starting point of development. This approach produced a pseudotime value for each cell that reflects their progression in maturation. Pseudotime correctly increased, as MSNs moved from early and apical progenitors towards mature MSNs (**Fig. 5A**). We further evaluated the expression of several neuronal maturation markers in relation to pseudotime to ensure correspondence with maturation state. As expected, we observed a decline in *NES* levels as pseudotime advanced (**Fig. 5B**). *ASCL1*, an intermediate progenitor marker, accurately demonstrated peak expression in the middle phases of fetal and C116 MSN development (**Fig. 5C**). However, developing HD72 MSNs did not exhibit increased *ASCL1* expression at any point in pseudotime. We also examined the expression of *DCX*, a marker of developing neurons previously identified to exhibit elevated expression in basal progenitors and newly born MSNs [31]. As expected, DCX expression increased along with pseudotime progression (**Fig. 5D**).

**Figure 5.**
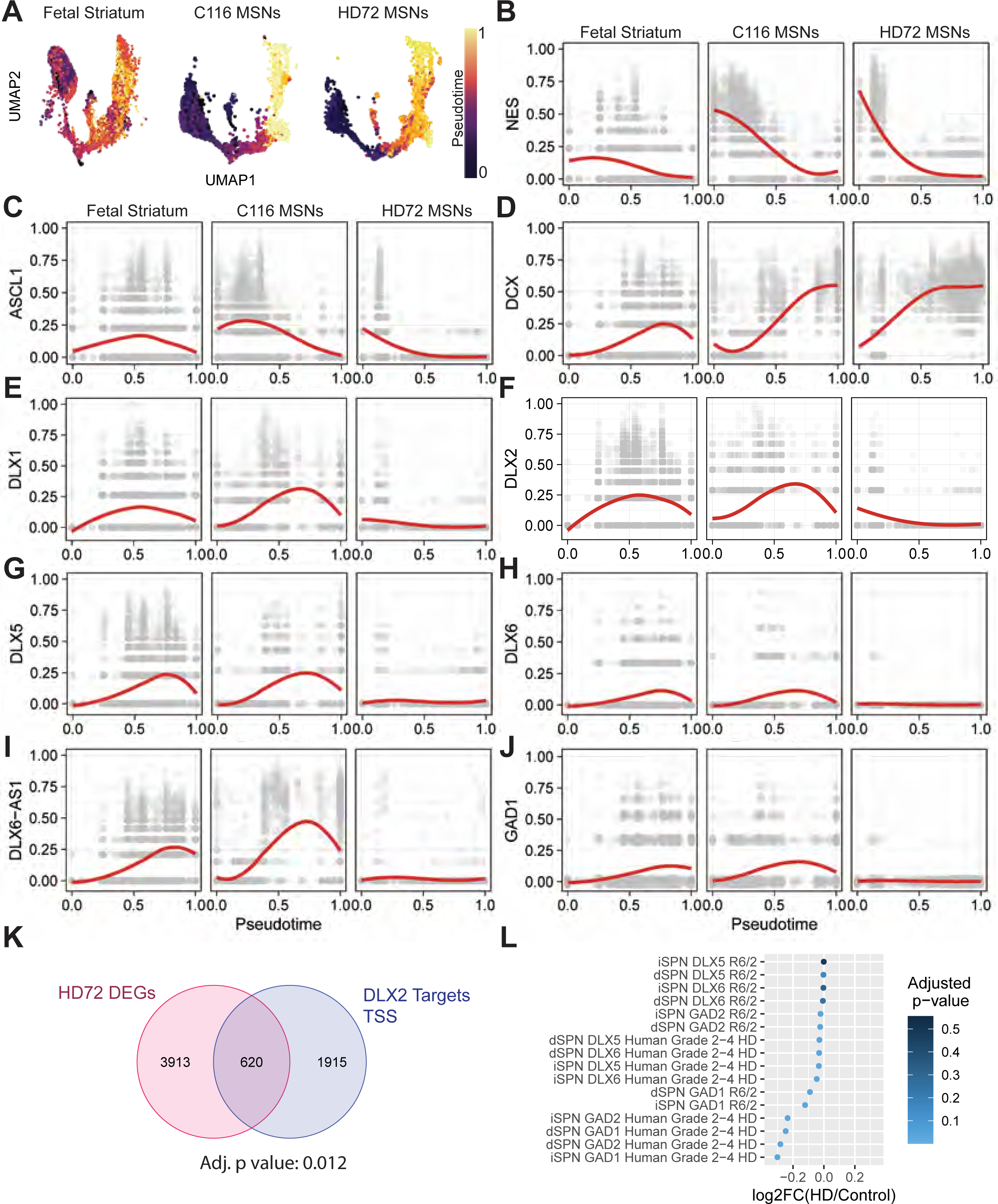
Developmental Dysregulation in Developing HD72 MSNs. **(A)** Pseudotime in fetal LGE, C116 MSNs and HD72 MSNs. Expression of neuronal maturation markers, NES **(B)**, ASCL1 **(C)** and DCX **(D)**. Expression of transcription factors DLX1 **(E)**, DLX2 **(F)**, DLX5 **(G)**, DLX6 **(H)**, DLX6-AS1 **(I)**, GAD1 **(J)** in human LGE and developing C116 and HD72 MSNs. **(K)** Comparison of DLX2 targets at transcriptional start sites with DEGs in HD72 MSNs. P values were calculated using Fisher’s exact tests followed by Benjamini-Hochberg correction for multiple tests. **(L)** Expression of DLX5, DLX6, GAD1 and GAD2 in iSPNs and dSPNs from caudate and putamen of HD patients and R6/2.

ASCL1 is a critical transcription factor that promotes neuronal differentiation [61]. Within the LGE, ASCL1 regulates the expression of DLX1 and DLX2, which are required for proper striatal development. [62-67]. The downregulation of *ASCL1* during HD MSN maturation raises concerns about the potential impact on its downstream targets. To investigate this, we assessed the expression of *DLX1/2* and other members of the *DLX* family as a function of pseudotime. While *DLX1/2/5/6/6-AS1* expression increased during the midst of fetal and C116 MSNs development, this upregulation failed to occur in HD72 MSNs (**Fig. 5E-I**).

In addition, we observed a similar pattern of downregulation for glutamate decarboxylase 1 (*GAD1*), a GABAergic marker that is regulated by the *DLX* TFs [68, 69] (**Fig. 5J**). To explore the broader impact of *DLX* TF dysregulation in HD72 MSNs, we examined whether other target genes were affected. Specifically, we compared the differentially expressed genes in developing HD72 MSNs with those identified as targets for *Dlx1*, *Dlx2*, and *Dlx5* during murine LGE development [69]. Our analysis revealed a significant enrichment of Dlx2 targets among the DEGs in developing HD72 MSNs, with 620 genes shared between the two sets (adjusted p-value: 0.017) (**Fig. 5K**). Furthermore, these 620 genes were bound by Dlx2 at their transcriptional start site in CHIP-seq experiments of developing mouse striatum [69].

To assess whether the dysregulation of *DLX* genes persisted into adulthood and disease onset, we analyzed the expression of *DLX* TFs, *GAD1*, and *GAD2* in a published scRNAseq dataset from the caudate and putamen of HD patients and striatum from the R6/2 [70]. *DLX1* and *DLX2* expression was sparse in iSPNs and dSPNs of human and mouse datasets and did not permit a relevant comparison of HD and control samples. *DLX5* and *DLX6* were downregulated in iSPNs and dSPNs of HD patients. *GAD1* and *GAD2* were also downregulated (**Fig. 5J**). In R6/2 iSPNs and dSPNs, we observed downregulation of *Dlx6*, *Gad1*, and *Gad2*, although *Dlx6* was only downregulated in R6/2 dSPNs and not in iSPNs (**Fig. 5J**).

In addition to pseudotime plots, we also compared the expression of *DLX1*, *DLX2*, *DLX5*, *DLX6*, *DLX6-AS1*, *GAD1* and *GAD2* in the various clusters corresponding to different stages of development. We found a lower proportion of cells expressing these genes in HD72 MSNs (**Supplemental Fig. 4A-G**). We also observed higher expression of cortical markers neurogenin 2 (*NEUROG2*), vesicular glutamate transporter (*SLC17A6/VGLUT2*), EBF transcription factor 3 (*EBF3*), and nescient helix-loop-helix 2 (*NHLH2*) in HD72 MSNs (**Supplemental Fig. 4H-K**). These markers suggest a partial loss of cell identity, as reported in other HD models [44].

### Predicted Developmental HD Modifiers Based on Transcriptional Data can Reverse HD Phenotypes

A promising strategy for identifying disease modifiers is to search for small molecules that induce gene expression patterns that are the opposite of those in the disease state. This approach has been successfully used in Alzheimer’s and Parkinson’s disease models [71, 72]. To demonstrate the feasibility of this approach in HD, we utilized the L1000CDS2 web application [73]. Leveraging the L1000 dataset, L1000CDS2 identified small molecules that produce an opposite or similar gene expression pattern to a given input.

We used DEGs from developing HD72 MSNs identified by bulk RNAseq and their log2-fold change as input to obtain 50 signatures of small molecules that cause changes opposite to those seen in HD (**Supplemental Table 6**). Several of the resulting small molecules (e.g., withaferin-a, celastrol, trichostatin A, vorinostat, and niclosamide), have beneficial effects in mouse and cellular HD models (**Fig. 6A**) [74-78]. We further categorized the predicted small molecules based on their canonical targets and identified two major classes that showed efficacy in HD models: histone deacetylase (HDAC) inhibitors and epidermal growth factor receptor (EGFR) inhibitors (**Fig. 6B**) (**Supplemental Table 6**) [75, 76, 79].

**Figure 6.**
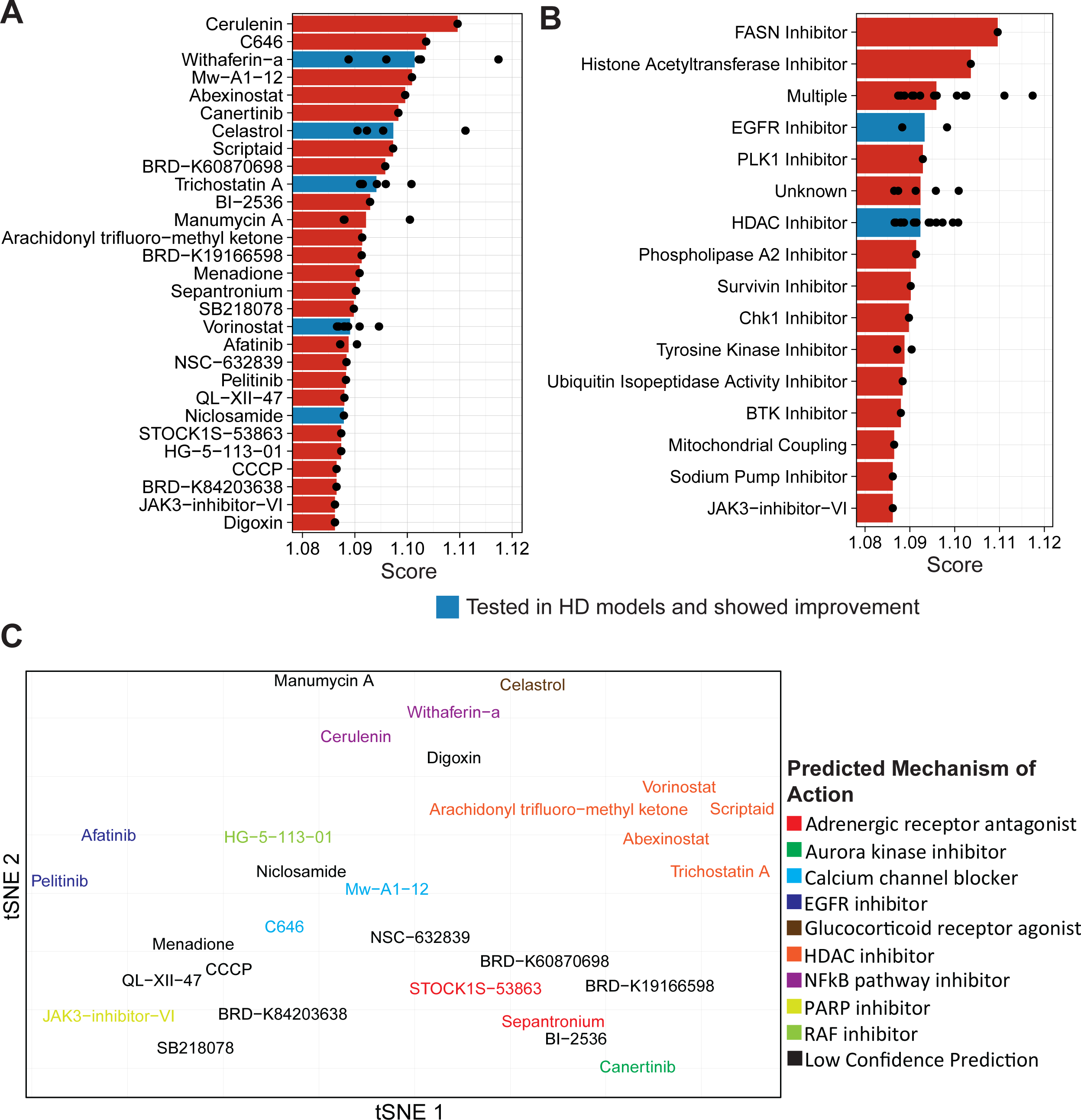
Small Molecules Predicted to Reverse HD Dysregulation. **(A)** List of small molecules predicted to reverse gene dysregulation in developing HD72 MSNs. Each dot represents an individual signature obtained from treatment of a cell line with the small molecule. **(B)** Canonical targets for predicted small molecules. **(C)** tSNE plot based on scores of predicted mechanisms of action for each small molecule. Molecules with more than a 40% confidence are color coded to its top mechanism of action.

To explore potential non-canonical mechanisms of action, we utilized the L1000FWD portal, a web application to predicted mechanisms of action (MOAs) based on the transcriptional profile produced by the small molecules (**Supplemental Table 6**) [80]. We used the probability scores that accompany each predicted MOA to generate a t-distributed stochastic neighbor embedding (tSNE) plot and colored the small molecules by their top predicted MOA (**Fig. 6C**). Small molecules without MOAs with probability scores above 40% were not labeled.

The top predicted MOA for Afatinib and Pelitinib correctly labeled them as EGFR inhibitors. Similarly, Vorinostat, Scriptaid, Abexinostat, and Trichostatin-A are known HDAC inhibitors that were correctly classified. While arachidonyl trifluoromethyl ketone is not typically acknowledged as an HDAC inhibitor, trifluoromethyl ketones indeed act as inhibitors for HDACs [81]. Manumycin A and Cerulenin were labeled as NF-κB inhibitors and both molecules disrupt in NF-κB signaling [82-85]. JAK3-inhibitor-VI was classified as a poly(ADP-ribose) polymerase 1 (PARP) inhibitor. Canertinib, although canonically recognized as an EGFR inhibitor, received high scores as an aurora kinase inhibitor, which has been linked to EGFR activity in cancer [86]. Additionally, STOCK1S-53863 and Sepantronium were predicted to be adrenergic receptor antagonists, and C646 and Mw-A1-12 were classified as calcium channel blockers. Finally, HG-5-113-01 was predicted to be a rapidly accelerated fibrosarcoma (RAF) inhibitor (**Fig. 6C**).

The presence of multiple validated small molecules in cell and mouse models of HD confirms the effectiveness of this approach in finding HD modifiers. To further investigate, the efficacy of an untested molecule, we chose to test Cerulenin’s ability to reverse phenotypes in developing HD72 MSNs. Among the predicted molecules, Cerulenin’s transcriptomic signature had one of the highest scores. Although it has not been tested in HD models, its mechanisms of action are similar to Withaferin A’s, which is beneficial in HD models [77]. Cerulenin’s transcriptional signature showed modulation of multiple members of the BDNF signaling pathway (**Supplemental Table 6**). BDNF treatment improves HD phenotypes, including a rescue in the expression of DARPP-32, a phenotype that our model of HD MSNs recapitulates at the protein and RNA level (**Fig. 1C, Supplemental Figure 3C**) [47, 56, 87, 88]. Thus, we treated developing HD72 MSNs with doses of Cerulenin of 31.25–2000 nM starting on day 11 of differentiation. After 9 days of treatment, DARPP-32 expression was assessed via immunofluorescence. Cerulenin caused a dose-dependent increase in the expression of DARPP-32, which plateaued at 250 nM (**Fig. 7A**). Upon treating C116 and HD72 developing MSNs with 250 nM Cerulenin, we observed an increase in DARPP-32 expression exclusively in HD72 MSNs, but no significant change was detected in C116 MSNs (**Fig. 7B, C**). We also tested the electrical activity of C116 and HD72 MSNs on a multi-electrode array (MEA) plate. C116 MSNs displayed a higher rate of active electrodes per minute (**Fig. 7D**), in line with previous reports showing decreased electrical activity in iPSC derived neurons [89, 90]. Treatment with 250nM Cerulenin was able to increase the rate of active electrodes per minute significantly.

**Figure 7.**
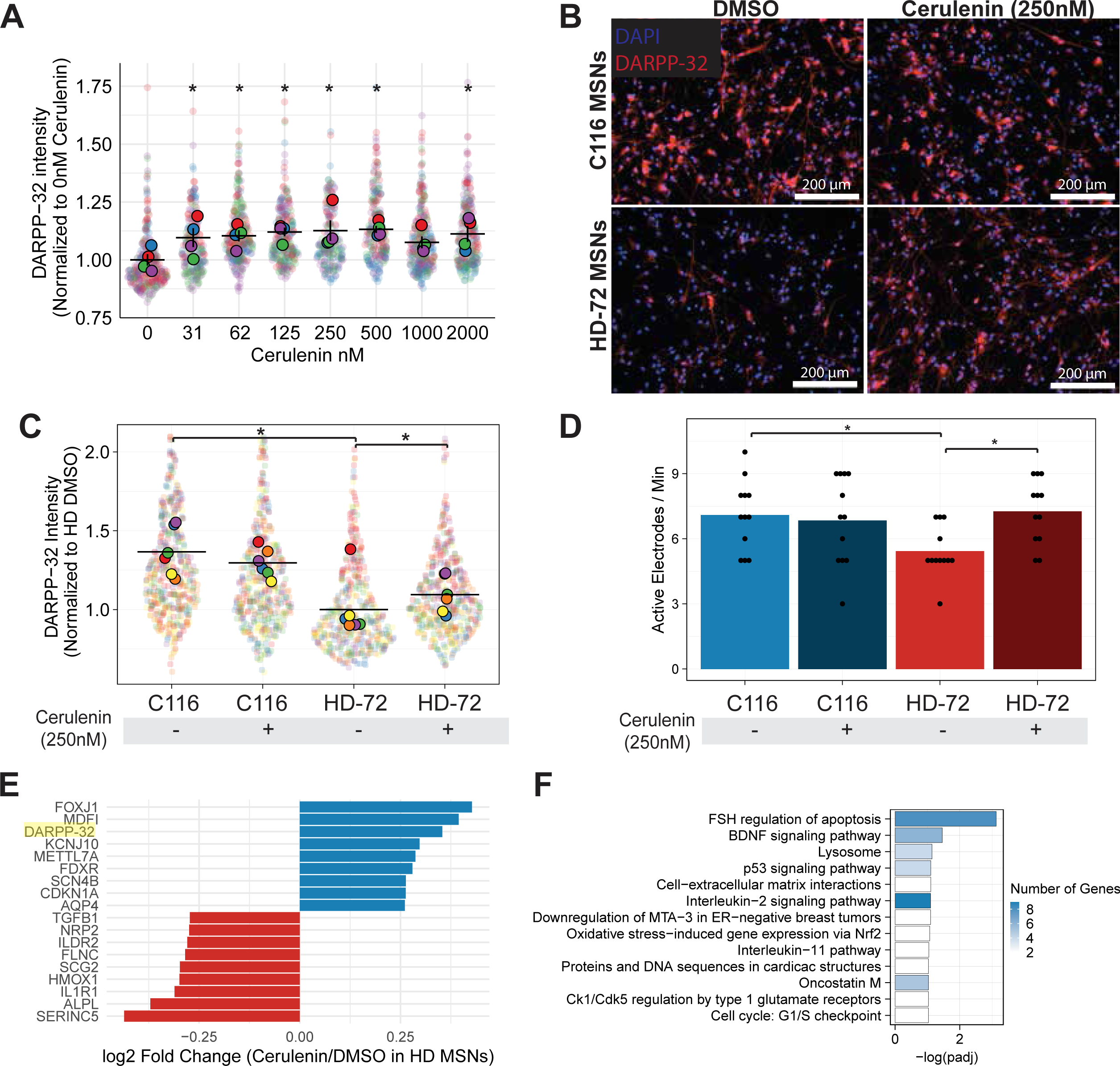
Effects of Cerulenin Treatment on HD72 MSNs. **(A)** Quantification of DARPP-32 levels from immunostaining of Cerulenin treated HD72 developing MSNs. P values were calculated using Dun’s test followed by Benjamini-Hochberg correction for the comparisons to HD DMSO. **(B)** Immunostaining of HD72 and C116 MSNs treated with 250nM Cerulenin. Scale bar: 200 μm. **(C)** Quantification of DARPP-32 in C116 and HD72 MSNs treated with 250nM Cerulenin. P values were calculated using pairwise Wilcoxon’s tests followed by Benjamini-Hochberg correction **(D)** Active electrodes/minute in cultures of C116, and HD72 MSNs treated with DMSO or 250nM Cerulenin. P values were calculated using pairwise Wilcoxon’s tests followed by Benjamini-Hochberg correction. **(E)** Top genes modulated in RNAseq of HD MSNs treated Cerulenin. **(F)** Dot plot showing pathways altered by Cerulenin treatment in HD72 MSNs.

RNAseq of HD72 and C116 MSNs treated with Cerulenin confirmed upregulation of DARPP-32 (*PPP1R1B*) (**Fig. 7E**) and showed changes in genes involved in FSH regulation of apoptosis, BDNF signaling pathway, lysosome, p53 singnaling pathway, and cell-extracellular matrix interactions among others (**Fig. 7F**) (**Supplemental Table 7**). In sum, these data suggest that modulation of these pathways by Cerulenin underlies its effects in HD72 MSNs and, warrants further in-depth studies. To identify perturbations that caused transcriptional effects similar to those caused by Cerulenin on HD72 MSNs, we used a SigCom LINCS signature search [91] with the gene expression signature obtained from HD72 MSNs treated with Cerulenin. The most similar chemical perturbation was caused by Whitaferin A, confirming their similarity (**Supplemental Table 7**). Despite Cerulenin’s canonical target being fatty acid synthase (FASN), the signature search did not find a similar signature between cells treated with FASN siRNA or a FASN CRISPR knockdown system and those treated with Cerulenin. This suggests that Cerulenin interacts with targets in addition to FASN as part of its mechanism of action in HD72 MSNs.

## DISCUSSION

Recent research has shed light on the impact of HD on neuronal development and its potential pathological consequences [7, 12, 28]. These findings suggest the possibility of new therapeutic approaches and underscore the need to gain a better understanding of how mHTT alters brain development and whether those changes sets the stage for the disorders observed later in life.

In this study, we utilized isogenic HD72 and corrected iPSCs-derived MSNs to investigate the effects of HD on the development of human MSNs as they differentiate from NSCs to mature MSNs. Consistent with previous studies [26], our results show that iPSCs-derived developing MSNs recapitulate multiple aspects of fetal striatum development. Specifically, we observed similar gene-expression patterns and cell-type compositions between the two systems, including early and intermediate progenitors and mature MSNs. However, we also noted that iPSC-derived early progenitors lacked expression of *TCF7L1* and *GSX2*, indicating a difference in the cellular identity at this stage between iPSC-derived early progenitors and the human LGE.

After establishing the degree to which our model faithfully recapitulates MSNs development, we investigated how the HD mutation affects this process. We observed abnormal neuronal maturation in HD72 developing MSNs. Specifically, premature maturation was indicated by an accelerated decline in *NES* expression and an increase in neuronal maturations markers. This finding agrees with previous studies that found accelerated maturation of human HD NSCs implanted in mice [22] and is consistent with the hypertrophy seen in child HD carriers [15].

These developmental dysregulations appear to be associated with the aberrant expression of factors that regulate neuronal differentiation. For example, *ASCL1*, a transcription factor that promotes neuronal differentiation [61], is upregulated in intermediate MSN progenitors, but its expression is significantly downregulated in HD72 intermediate progenitors. However, even in the absence of *ASCL1*, mature HD72 MSNs express neuronal maturation markers, suggesting that an atypical maturation process is responsible for their development. Notably, *NEUROG2*, which is also involved in neuronal maturation, is upregulated in HD72 intermediate progenitors, indicating an abnormal compensatory MSN maturation program due to the lack of ASCL1. This is notable since cases of *ASCL1* and *NGN2* with a compensatory relationship in the central nervous system have been documented [92].

We found dysregulation of *ASCL1* correlated with dysregulation of DLX transcription factors, that are regulatory targets of ASCL1 and necessary for proper striatum development [66, 67]. A lack of DLX expression during development leads to ectopic expression of cortical markers in the striatum and to a reduction of genes essential for striatal function and development [67]. In agreement with these studies, we observed that HD72 mature MSNs have lower levels of key MSN genes (e.g., *GAD1* and *GAD2*) and ectopic expression of genes normally expressed in the cortex (e.g., *SLC17A6*, *EBF3* and *NHLH2*). In addition, we found that multiple other genes known to be targets of *DLX2* during striatum development [69] are dysregulated in developing HD72 MSNs. Moreover, an examination of published datasets from both Grade 2–4 HD patient MSNs and the R6/2 mouse model [70] indicated reductions in *DLX5*, *DLX6*, *GAD1*, and *GAD2* expression, suggesting that some of the dysregulation during early development could continue or recur during disease onset.

Our analysis also revealed changes in multiple signaling pathways related to HD pathology. Bulk RNAseq of NSCs and MSNs showed enrichment for pathways known to play roles in HD, such as BDNF signaling, TGF-β signaling, axon guidance, synaptogenesis, and NRF2-mediated oxidative stress response [46, 53, 93-95]. Dysregulation of the BDNF pathway has been linked to disturbances in neuronal circuits during development [96]. The disruption of BDNF signaling and the abnormal maturation of HD MSNs could potentially lead to defects in neuronal circuitry within the striatum, with pathogenic consequences later on. This phenomenon is observed in the cortex of HD mice, where defects in neuronal circuitry contribute to the disease’s development [12]. Early correction of these changes offers protection against the disease’s progression later in life [12].

scRNAseq of developing MSNs allowed us to examine the dysregulation of signaling pathways as HD72 MSNs matured. For instance, as HD72 MSNs mature, the NRF2-mediated oxidative stress response is predicted to increase, suggesting that the HD mutation leads to a greater increase in oxidative stress with advancing MSN maturation. Conversely, the ERK/MAPK signaling pathway follows an opposing pattern, with high levels of activation in HD72 progenitors that decline as MSNs mature. ERK activation is altered by HD, and it’s activation is protective in multiple models [55]. Thus, immature HD progenitors might use it as a protective mechanism that is lost as MSNs mature. These findings suggest that dysregulation of these pathways in HD has its origins during MSN development and highlight the potential for early intervention and influence disease progression.

To identify HD modifiers from transcriptomic data, we searched for small molecules that induce gene expression changes opposite to those in developing HD72 MSNs. Our predictions identified several that improve HD pathology. Notably, HDAC inhibitors, Celastrol, and Withaferin-A were the most robust interventions and, displayed multiple gene expression signatures predicted to reverse HD dysregulation. These molecules also demonstrated efficacy in HD models [75-78].

To assess the effects of a predicted small molecule untested in HD, we treated developing HD72 MSNs with Cerulenin. Cerulenin is a fatty acid synthesis inhibitor, and modulates NF-kB and eIF2α signaling [83, 97]. Our predictions indicated that it would affect components of the BDNF signaling pathway which is linked to the downregulation of DARPP-32 expression [98], a hallmark of HD [56, 99-101]. HD72 MSNs treated through half of the differentiation period with Cerulenin displayed a dose-dependent increase in DARPP-32 levels that plateaued at 250 nM. In addition, levels of electrical activity were lower in iPSCs-derived HD72 MSNs. The low level of electrical activity in HD MSNs has previously been reported [89, 90]. We find Cerulenin rescues this but not to control levels. Cerulenin did not show similar effects on C116 MSNs, suggesting that the mechanism is specific for HD. RNAseq analysis of Cerulenin-treated HD72 MSNs revealed changes in p53 signaling and other pathways associated with HD pathology [102]. However, levels of DLX transcription factors or their targets were not improved. Interestingly, reduction of HDAC4 increases levels of *DLX1/2/5/6* in the R6/2 model [103], suggesting HDAC inhibitors might rescue those deficits.

In summary, we examined the impact of HD on MSNs development using iPSC and isogenic controls, uncovering dysregulation of key genes for proper maturation. We discovered aberrant expression of ASCL1 and DLX transcription factors and dysregulation of HD-related pathways, such as NRF2-mediated oxidative stress response and ERK/MAPK signaling as MSNs mature. We also provide proof of concept for identification of HD modifiers from transcriptional data by showing a partial rescue of DARPP-32 levels and electrical activity in HD MSNs after treatment with Cerulenin as well as predicting multiple small molecules already confirmed to have beneficial effects in HD models.

## EXPERIMENTAL PROCEDURES

### NSC and developing MSN Differentiation

NSCs and MSNs were differentiated as described in [38]. Briefly, HD72 and C116 iPSCs were turned into cell aggregates and driven towards a neuroepithelial fate that produced neural rosettes. Neural rosettes were manually picked and dissociated into single cells to produce NSCs. NSCs were then expanded in Neurobasal medium (Thermo Fisher Scientific, 21103049) supplemented with B27-supplement 1 X (Thermo Fisher Scientific, 17504001), GlutaMAX 1 X (Thermo Fisher Scientific, 35050061), 10 ng/mL leukemia inhibitory factor (PeproTech, 300-05), and 100 U/mL penicillin-streptomycin.

To produce developing MSNs, NSCs were cultured in Synaptojuice A medium supplemented with 25 ng/mL of Activin A (PeproTech, AF-120-14E) while changing half of the medium every other day. After 7 days, the cells were switched to Synaptojuice B medium supplemented with 25 ng/mL of Activin A for 14 days. The cells were harvested or used for assays on day 20 of differentiation.

### Cerulenin Treatment

2.23 mL of DMSO (Sigma, D2650-5X10mL) were added to 1 mg of Cerulenin (Cayman Technical, No. 10005647) to make a stock solution of 2 mM. Cerulenin was diluted in Synaptojuice B to the desired concentration and added to the cells starting at day 11 of differentiation and every time the medium was changed until the cells were harvested. The cells were used for immunocytochemistry or RNAseq on day 20 of differentiation. For use with MEA plates, the cells were differentiated for 35 days and cerulenin was added every time the medium were changed after day 11 of differentiation.

### Multielectrode Array Measurements

Cytoview MEA plates (Axion Biosystems, M384-tMEA-24W) were coated with 1% poly(ethyleneimine) solution (Sigma-Aldrich, 03880-500mL) overnight at 37 °C. The next day, each well was rinsed with cell-culture grade water 3 times and allowed to dry for 2 hours in a cell-culture hood. Once the plates were dry, they were coated with 0.25μg/mL laminin (Sigma-Aldrich, L2020-1MG) in DPBS for 3 hours at 37 °C. NSCs were seeded at a density of 49,586 cells/mm^2^, and they were differentiated as described above until day 20. After day 20, half the medium was changed every 7 days, and readings were taken at day 35 of differentiation. The cells were placed inside a Maestro Edge (Axion Biosystems) instrument with temperature set at 37 °C and 5% CO_2_. Recordings were taken for 5 minutes, and the last minute was utilized for analysis. Electrical activity was assessed using a spike detector with adaptive thresholding set to 4.5 standard deviations. Metrics for electrical activity were generated using Axis Navigator and Neural Metric Tool software (Axion Biosystems).

### Immunofluorescence

Cells were rinsed with PBS and then fixed in 4% paraformaldehyde in PBS for 12 minutes at room temperature. Fixation was followed by three washes of PBS for 5 minutes each. Fixed cells were blocked using blocking buffer containing 0.1% Triton-x-100 (Thermo Fisher Scientific, 28313), 4% normal donkey serum (Jackson Immuno Research, 017-000-121) in PBS for 1 hour. Fixed cells were then incubated overnight with primary antibodies diluted 1:100 in blocking buffer. Cells were washed three times with PBS containing 0.1% Triton-x-100 and incubated with secondary antibodies diluted 1:500 in blocking buffer for 2 hours. Cells were washed 3 times for 5 minutes with PBS containing 0.1% Triton-x-100. The cells were finally placed in PBS and imaged utilizing a Cytation 5 (Biotek). The antibodies used were mouse anti-DARPP-32 antibody (Santa Cruz Biotechnologies, sc-271111) paired with donkey-anti-mouse Alexa-647 (Thermofisher, A-31571), Nestin (abcam, ab92391) paired with donkey-anti-rabbit Alexa-594 (Thermofisher, A-21207), Nestin (Santa Cruz Biotechnologies, sc-33677) paired with goat-anti-mouse Alexa-488 (Thermofisher, A11006), and SOX2 (Cell Signaling, 14962S) paired with goat-anti-rabbit Alexa 555 (Thermofisher, A21428).

### Bulk RNAseq Processing and Analysis

RNA was extracted from MSNs utilizing an RNA extraction kit (Bioline, BIO-52073). RNA library preparation was prepared at the UC Davis Genomics Core or Novogene, utilizing a poly-A library prep. Resulting FASTA files were then aligned utilizing STAR 2.7.10a to the GRCh38 primary assembly genome reference. Features were counted with the summarizeoverlaps function part of the GenomicAlignments R package [104]. Differential expression was performed with DESeq2 [105], and volcano plots were done with ggplot2. DEGs (adjusted p value < 0.01, mean base > 10 and absolute log2 fold-change > 0.1) were used as input for IPA version 76765844 to obtain prediction of canonical pathways. The p-values for enrichment analysis were corrected using the Benjamini-Hochberg correction. Enrichr was used to identify enrichment from the bioplanet database [106, 107]. To compare the DEGs in HD MSNs with those in other HD models, we obtained DEGs from [39-43] and performed a Fisher’s exact test to compare HD MSN DEGs with each of the HD models and adjusted p-values using Benjamini-Hochberg correction. Leafcutter 0.2.9 was used to detect differentially spliced genes in C116 and HD MSNs.

### scRNAseq Processing

NSCs and MSNs were cultured in six well plates coated with 50 µg/mL of Matrigel (Corning) overnight. Cells were treated with Accutase (Thermo Fisher, A1110501) for 5 minutes, and the cells were centrifuged and resuspended in DPBS. Cells were encapsulated, barcoded and transformed into libraries utilizing a Chromium Next GEM Single Cell 3ʹ Kit v3.1. Resulting libraries were sent for sequencing to the UC Davis genomics core and sequenced in a Novaseq 6000 lane. FASTQ files were aligned to the genome and used to generate gene counts utilizing cell ranger 6.0 and the GRCh38 reference genome. The Seurat workflow was used for filtering, normalization, scaling, PCA, UMAP generation, and clustering. The Seurat objects and C116 and HD datasets were integrated for visualization using Harmony [108]. To identify differentially expressed genes, we utilized the function FindMarkers and used MAST as the differential expression test. The results were used as input for IPA, filtering for genes with an adjusted p-value under 0.05 and an absolute log2-fold change larger than 0.25. The set of detected genes was used as background for IPA and p-values for enrichment were adjusted using the Benjamini-Hochberg correction.

### Comparison of MSN and LGE scRNAseq data

We obtained the raw data from the LGE dataset, which was originally published in Bocchi et al [31], by downloading it from the ArrayExpress database (www.ebi.ac.uk/arrayexpress/) under accession number E-MTAB-8894. FASTQ were processed as described above. The LGE dataset was down sampled using the subset function from the Seurat package so that it would match the number of cells found in the MSN datasets. Clusters of cells not positive for MEIS2, indicating a non LGE lineage, were removed. C116 and HD developing MSNs were integrated to the LGE dataset using Harmony [108]. Trajectory inference and pseudotime were produced with VIA [60].

### Drug Prediction and MOA Inference

DEGs identified in developing HD and C116 MSNs via Bulk RNAseq were used as input for the L1000CDS2 web application. The log2-fold change of the genes was used to indicate magnitude. Whether the predicted small molecules had already been used in HD models and their canonical targets were determined doing a manual literature search on PubMed. The predicted MOAs for each small molecule were obtained from the L1000FWD web application. The probability for each MOA in each small molecule signature was used to generate a tSNE plot using the function Rtsne from the Rtsne package. We manually colored small molecules in the tSNE that had a top MOA prediction with more than a 40% probability.

### Quantitative PCR

RNA extraction was carried out using the RNeasy Plus Mini Kit (QIAGEN, 74034), followed by its conversion into cDNA with the SensiFAST cDNA Synthesis Kit (Bioline, BIO-65053). qPCR was conducted using the SensiFAST Probe No-ROX Kit (Bioline, BIO-86005), UPL probes (Roche) and primers listed in **Supplementary Table 8**, on the LightCycler 480 system (Roche). For quantification purposes, the threshold cycle (Cp) for each amplification was identified using the 2nd derivative analysis provided by the LightCycler 480 software. Subsequently, the 2-ΔΔCp method was applied to calculate the relative expression levels of individual genes, which were normalized against the housekeeping gene β-actin (ACTB).

## SUPPLEMENTAL FIGURE LEGENDS

**Supplemental Figure 1. Comparison with proteomics and differential splicing analysis. (A)** Plot showing correlation between log2-fold change of genes in HD72 NSCs and HD72 MSNs. **(B)** Plot showing correlation log2-fold changes in RNA from HD72 MSNs versus changes in proteins. **(C)** Venn diagram showing overlap of differentially spliced genes between HD72 MSNs, human postmortem striatum and R6/1 striatum. **(D)** Enrichment for genes differentially spliced in MSNs that are shared with human HD striatum or R6/1 striatum.

**Supplemental Figure 2. scRNAseq of C116 and HD72 NSCs. (A)** C116 and HD72 NSCs UMAP plots divided into four different clusters. **(B)** Heatmap showing differentially expressed genes between the identified clusters. **(C)** Expression of Nestin in C116 and HD72 NSCs. **(D)** Violin plots showing levels of top downregulated (UBB, RBP1 and CRYBB1) and upregulated (OCIAD2, IGFBP5 and S100B) genes in HD72 NSCs clusters.

**Supplemental Figure 3. Proportion of Early Progenitors in Developing MSNs. (A)** Nestin labeled C116 and HD72 developing MSNs. Scale bar: 100 μm. **(B)** Distribution of Nestin-C116 and HD72 in developing MSNs. P value was calculated using the Kolmogorov-Smirnov test. **(B)** Levels of DARPP-32 RNA in C116 and HD72 MSNs measured with qPCR. P value was calculated with a t test.

**Supplemental Figure 4. Expression of DLX Genes and Targets in Different Clusters in C116 and HD72 Developing MSNs.** Violin plots showing expression of DLX1 **(A)**, DLX2 **(B)**, DLX5 **(C)**, DLX6 **(D)**, DLX6-AS1 **(E)**, GAD1 **(F)**, GAD2 **(G)**, NEUROG2 **(H)**, SLC17A6 **(I)**, EBF3 **(J)** NHLH2 (**K**) in C116 and HD early progenitors, intermediate progenitors and mature MSNs.

**Supplemental Table 1**. DEGs in HD72 and C116 NSCs. IPA on canonical pathways predicted to be dysregulated in NSCs. Bioplanet enrichment analysis for HD72 NSCs.

**Supplemental Table 2**. DEGs in HD72 and C116 developing MSNs. IPA on canonical pathways predicted to be dysregulated in MSNs. Bioplanet enrichment analysis for HD72 developing MSNs. Comparison of DEGs found in HD72 developing MSNs and other HD models.

**Supplemental Table 3**. Differentially spliced genes in NSCs and developing MSNs. Enrichment analysis of shared differentially spliced genes in HD72 developing MSNs, R6/1 striatum, and HD patient striatum.

**Supplemental Table 4**. Single cell DEGs in HD72 NSCs, early progenitors, intermediate progenitors and mature MSNs. IPA of DEGs in early progenitors, intermediate progenitors and mature MSNs. IPA comparison analysis for canonical pathways dysregulated in early progenitors, intermediate progenitors and mature MSNs.

**Supplemental Table 5**. Comparison of DLX targets during striatum development with DEGs in HD72 developing MSNs.

**Supplemental Table 6**. Small molecules predicted to reverse transcriptional dysregulation in HD72-developing MSNs. Additional information on predicted small molecules and predicted mechanisms of action.

**Supplemental Table 7**. DEGs found after Cerulenin treatment in HD72-developing MSNs. IPA of DEGs found after Cerulenin treatment. Signatures of other small molecules with similarities to Cerulenin.

**Supplemental Table 8**. qPCR primers and probes used.

## Abbreviations

ASCL1: achaete-scute homolog 1; family BHLH transcription factor 1
BCL11B: BCL11 transcription factor B
BDNF: brain derived neurotrophic factor
CAG: cytosine adenine guanine
CALB1: Calbindin 1
CALB2: Calbindin 2
CLEAR: coordinated lysosomal expression and regulation
CNS: central nervous system
CTIP2: BCL11 transcription factor B
DARPP-32: dopamine and cAMP-regulated phosphoprotein of 32 kDa
DCX: Doublecortin
DEGs: differentially expressed genes
DLX: distal-less homeobox
DLX6-AS1: DLX6 antisense RNA 1
DRD1: dopamine receptor D1
DRD2: dopamine receptor D2
EBF1: early B cell factor-1
EBF3: early B-cell factor 3
EGFR: epidermal growth factor receptor
ERK: rxtracellular signal-regulated kinases
FASN: fatty acid synthase
FOXP1: forkhead box P1
GABA: gamma aminobutyric acid
GAD1: glutamate decarboxylase 1
GAD2: glutamate decarboxylase 2
GPC1: glypican 1
GPM6B: glycoprotein M6B
HD: Huntington’s disease
HDAC: histon deacetylase inhibitor
HES6: hes Family BHLH transcription factor 6
HTT: Huntingtin
IGFBP5: insulin like growth factor binding protein 5
IGFBP7: insulin-like growth factor binding protein 7
ILK: integrin linked kinase
IPA: ingenuity pathway analysis
iPSCs: induced pluripotent stem cells
LGE: lateral ganglionic eminence
MAP2: microtubule associated protein 2
MAPK: mitogen-activated protein kinase
MEA: multi electrode array
MEIS2: meis homeobox 2
METRN: Meteorin
MOR1-6TM: opioid receptor Mu 1
mHTT: mutant Huntingtin
MSNs: medium spiny neurons
MSX1: MSH homeobox 1
NEUROG2: Neurogenin 2
NES: Nestin
NHLH2: nescient helix-loop-helix 2
NGN2: neurogenin 2
NRF2: nuclear Factor erythroid 2-related factor 2
NR4A1: nuclear receptor subfamily 4 group A member 1
NSCs: neuronal stem cells
OCIAD2: OCIA domain containing 2
PARPP: poly(ADP-ribose) polymerase 1
PBS: phosphate buffered saline
PCA: principal component analysis
PTEN: phosphatase and tensin homolog
polyQ: polyglutamine
RAF: rapidly accelerated fibrosarcoma
RBP1: retinol binding protein 1
RHOGDI: Rho GDP dissociation inhibitor alpha
RHO: Ras homolog
RNA: ribonucleic acid
RNAseq: RNA sequencing
scRNAseq: single cell RNA sequencing
SLC17A6: solute carrier family 17 member 6
SP9: Sp9 transcription factor
TCEAL7: transcription elongation factor A like 7
TCF7L1: transcription factor 7-like 1
TGF-β: transforming growth factor beta
TTYH1: tweety family member 1
UBB: Ubiquitin B
VIM: Vimentin

## REFERENCES

1. Hedreen, J.C., et al., Neuronal loss in layers V and VI of cerebral cortex in Huntington’s disease. Neurosci Lett, 1991. 133(2): p. 257–61.

2. Cudkowicz, M. and N.W. Kowall, Degeneration of pyramidal projection neurons in Huntington’s disease cortex. Ann Neurol, 1990. 27(2): p. 200–4.

3. Group, H.s.D.C.R., A novel gene containing a trinucleotide repeat that is expanded and unstable on Huntington’s disease chromosomes.. Cell, 1993. 72(6): p. 971–83.

4. Barnat, M., et al., Huntington’s disease alters human neurodevelopment. Science, 2020. 369(6505): p. 787-793.

5. Molero, A.E., et al., Impairment of developmental stem cell-mediated striatal neurogenesis and pluripotency genes in a knock-in model of Huntington’s disease. Proceedings of the National Academy of Sciences, 2009. 106(51): p. 21900–21905.

6. Ring, K.L., et al., Genomic Analysis Reveals Disruption of Striatal Neuronal Development and Therapeutic Targets in Human Huntington’s Disease Neural Stem Cells. Stem Cell Reports, 2015. 5(6): p. 1023–1038.

7. Molero, A.E., et al., Selective expression of mutant huntingtin during development recapitulates characteristic features of Huntington’s disease. Proc Natl Acad Sci U S A, 2016. 113(20): p. 5736–41.

8. Zhang, N., et al., iPSC-based drug screening for Huntington’s disease. Brain Res, 2016. 1638(Pt A): p. 42–56.

9. Cirnaru, M.D., et al., Nuclear Receptor Nr4a1 Regulates Striatal Striosome Development and Dopamine D(1) Receptor Signaling. eNeuro, 2019. 6(5).

10. Mehler, M.F., et al., Loss-of-Huntingtin in Medial and Lateral Ganglionic Lineages Differentially Disrupts Regional Interneuron and Projection Neuron Subtypes and Promotes Huntington’s Disease-Associated Behavioral, Cellular, and Pathological Hallmarks. The Journal of Neuroscience, 2019. 39(10): p. 1892–1909.

11. Cirnaru, M.-D., et al., Unbiased identification of novel transcription factors in striatal compartmentation and striosome maturation. eLife, 2021. 10.

12. Braz, B.Y., et al., Treating early postnatal circuit defect delays Huntington’s disease onset and pathology in mice. Science, 2022. 377(6613): p. eabq5011.

13. Kim, H., et al., A pathogenic proteolysis–resistant huntingtin isoform induced by an antisense oligonucleotide maintains huntingtin function. JCI Insight, 2022. 7(17).

14. Rodríguez-Urgellés, E., et al., Postnatal Foxp2 regulates early psychiatric-like phenotypes and associated molecular alterations in the R6/1 transgenic mouse model of Huntington’s disease. Neurobiology of disease., 2022. 173: p. 105854.

15. van der Plas, E., et al., Abnormal brain development in child and adolescent carriers of mutant huntingtin. Neurology, 2019. 93(10): p. e1021–e1030.

16. Scahill, R.I., et al., Biological and clinical characteristics of gene carriers far from predicted onset in the Huntington’s disease Young Adult Study (HD-YAS): a cross-sectional analysis. Lancet Neurol, 2020. 19(6): p. 502–512.

17. Humbert, S., Is Huntington disease a developmental disorder? EMBO Rep, 2010. 11(12): p. 899.

18. Godin, J.D., et al., Huntingtin is required for mitotic spindle orientation and mammalian neurogenesis. Neuron, 2010. 67(3): p. 392–406.

19. Wiatr, K., et al., Huntington Disease as a Neurodevelopmental Disorder and Early Signs of the Disease in Stem Cells. Mol Neurobiol, 2018. 55(4): p. 3351–3371.

20. van der Plas, E., J.L. Schultz, and P.C. Nopoulos, The Neurodevelopmental Hypothesis of Huntington’s Disease. J Huntingtons Dis, 2020. 9(3): p. 217–229.

21. Smith-Geater, C., et al., Aberrant Development Corrected in Adult-Onset Huntington’s Disease iPSC-Derived Neuronal Cultures via WNT Signaling Modulation. Stem Cell Reports, 2020. 14(3): p. 406–419.

22. Miguez, A., et al., &lt;em&gt;In vivo&lt;/em&gt; progressive degeneration of Huntington’s disease patient-derived neurons reveals human-specific pathological phenotypes. bioRxiv, 2020: p. 2020.10.21.347062.

23. An, M.C., et al., Genetic correction of Huntington’s disease phenotypes in induced pluripotent stem cells. Cell Stem Cell, 2012. 11(2): p. 253–63.

24. Zhang, N., et al., Characterization of Human Huntington’s Disease Cell Model from Induced Pluripotent Stem Cells. PLoS Curr, 2010. 2: p. yfRRN1193.

25. Naphade, S., K.T. Tshilenge, and L.M. Ellerby, Modeling Polyglutamine Expansion Diseases with Induced Pluripotent Stem Cells. Neurotherapeutics, 2019. 16(4): p. 979–998.

26. Conforti, P., et al., In vitro-derived medium spiny neurons recapitulate human striatal development and complexity at single-cell resolution. Cell Rep Methods, 2022. 2(12): p. 100367.

27. Consortium, H.D.i., Induced pluripotent stem cells from patients with Huntington’s disease show CAG-repeat-expansion-associated phenotypes. Cell Stem Cell, 2012. 11(2): p. 264–78.

28. Song, S., et al., Postnatal Conditional Deletion of Bcl11b in Striatal Projection Neurons Mimics the Transcriptional Signature of Huntington’s Disease. Biomedicines, 2022. 10(10).

29. Arteaga-Bracho, E.E., et al., Postnatal and adult consequences of loss of huntingtin during development: Implications for Huntington’s disease. Neurobiol Dis, 2016. 96: p. 144–155.

30. Macdonald, M., A novel gene containing a trinucleotide repeat that is expanded and unstable on Huntington’s disease chromosomes. Cell, 1993. 72(6): p. 971–983.

31. Bocchi, V.D., et al., The coding and long noncoding single-cell atlas of the developing human fetal striatum. Science, 2021. 372(6542).

32. Tshilenge, K.-T., et al., Proteomic Analysis of Huntington’s Disease Medium Spiny Neurons Identifies Alterations in Lipid Droplets. bioRxiv, 2022: p. 2022.05.11.491152.

33. Lendahl, U., L.B. Zimmerman, and R.D. McKay, CNS stem cells express a new class of intermediate filament protein. Cell, 1990. 60(4): p. 585–95.

34. Kemp, P.J., et al., Improving and accelerating the differentiation and functional maturation of human stem cell-derived neurons: role of extracellular calcium and GABA. J Physiol, 2016. 594(22): p. 6583–6594.

35. Djousse, L., et al., Evidence for a modifier of onset age in Huntington disease linked to the HD gene in 4p16. Neurogenetics, 2004. 5(2): p. 109–14.

36. Benatar, T., et al., IGFBP7 reduces breast tumor growth by induction of senescence and apoptosis pathways. Breast Cancer Res Treat, 2012. 133(2): p. 563–73.

37. Voisin, J., et al., FOXO3 targets are reprogrammed as Huntington’s disease neural cells and striatal neurons face senescence with p16 ^INK4a^ increase. Aging Cell, 2020. 19(11).

38. Tshilenge, K.-T., et al., Proteomic Analysis of Huntington’s Disease Medium Spiny Neurons Identifies Alterations in Lipid Droplets. Molecular & cellular proteomics., 2023: p. 100534.

39. Gu, X., et al., N17 Modifies Mutant Huntingtin Nuclear Pathogenesis and Severity of Disease in HD BAC Transgenic Mice. Neuron, 2015. 85(4): p. 726–741.

40. Langfelder, P., et al., Integrated genomics and proteomics define huntingtin CAG length–dependent networks in mice. Nature Neuroscience, 2016. 19(4): p. 623–633.

41. Wertz, M.H., et al., Genome-wide In Vivo CNS Screening Identifies Genes that Modify CNS Neuronal Survival and mHTT Toxicity. Neuron, 2020. 106(1): p. 76–89 e8.

42. Nekrasov, E.D., et al., Manifestation of Huntington’s disease pathology in human induced pluripotent stem cell-derived neurons. Molecular Neurodegeneration, 2016. 11(1).

43. Hodges, A., et al., Regional and cellular gene expression changes in human Huntington’s disease brain. Human molecular genetics., 2006. 15(6): p. 965–977.

44. Malaiya, S., et al., Single-Nucleus RNA-Seq Reveals Dysregulation of Striatal Cell Identity Due to Huntington’s Disease Mutations. J Neurosci, 2021. 41(25): p. 5534–5552.

45. Battaglia, G., et al., Early defect of transforming growth factor beta1 formation in Huntington’s disease. J Cell Mol Med, 2011. 15(3): p. 555–71.

46. Capizzi, M., et al., Developmental defects in Huntington’s disease show that axonal growth and microtubule reorganization require NUMA1. Neuron, 2022. 110(1): p. 36–50.e5.

47. Zuccato, C., et al., Progressive loss of BDNF in a mouse model of Huntington’s disease and rescue by BDNF delivery. Pharmacol Res, 2005. 52(2): p. 133–9.

48. Zuccato, C. and E. Cattaneo, Role of brain-derived neurotrophic factor in Huntington’s disease. Progress in neurobiology, 2007. 81(5-6): p. 294–330.

49. Li, Y.I., et al., Annotation-free quantification of RNA splicing using LeafCutter. Nat Genet, 2018. 50(1): p. 151–158.

50. Elorza, A., et al., Huntington’s disease-specific mis-splicing unveils key effector genes and altered splicing factors. Brain, 2021. 144(7): p. 2009–2023.

51. Kung, V.W., et al., Dopamine-dependent long term potentiation in the dorsal striatum is reduced in the R6/2 mouse model of Huntington’s disease. Neuroscience, 2007. 146(4): p. 1571–80.

52. Singer, E., et al., The Novel Alpha-2 Adrenoceptor Inhibitor Beditin Reduces Cytotoxicity and Huntingtin Aggregates in Cell Models of Huntington’s Disease. Pharmaceuticals, 2021. 14(3): p. 257.

53. Tucci, P., et al., Nrf2 Pathway in Huntington’s Disease (HD): What Is Its Role? International Journal of Molecular Sciences, 2022. 23(23): p. 15272.

54. Di Pardo, A., et al., Defective Sphingosine-1-phosphate metabolism is a druggable target in Huntington’s disease. Scientific Reports, 2017. 7(1).

55. Bodai, L. and J.L. Marsh, A novel target for Huntington’s disease: ERK at the crossroads of signaling. BioEssays., 2012. 34(2): p. 142–148.

56. van Dellen, A., et al., N-Acetylaspartate and DARPP-32 levels decrease in the corpus striatum of Huntington’s disease mice. Neuroreport, 2000. 11(17): p. 3751–7.

57. Feng, J., et al., The Evf-2 noncoding RNA is transcribed from the Dlx-5/6 ultraconserved region and functions as a Dlx-2 transcriptional coactivator. Genes & Development, 2006. 20(11): p. 1470-1484.

58. Han, J., et al., OCIAD2 activates γ-secretase to enhance amyloid β production by interacting with nicastrin. Cellular and Molecular Life Sciences, 2014. 71(13): p. 2561–2576.

59. Sanada, F., et al., IGF Binding Protein-5 Induces Cell Senescence. Frontiers in endocrinology., 2018. 9.

60. Stassen, S.V., et al., Generalized and scalable trajectory inference in single-cell omics data with VIA. Nat Commun, 2021. 12(1): p. 5528.

61. Imayoshi, I., et al., Oscillatory Control of Factors Determining Multipotency and Fate in Mouse Neural Progenitors. Science., 2013. 342(6163): p. 1203-1208.

62. Anderson, S.A., et al., Mutations of the Homeobox Genes Dlx-1 and Dlx-2 Disrupt the Striatal Subventricular Zone and Differentiation of Late Born Striatal Neurons. Neuron., 1997. 19(1): p. 27–37.

63. Guo, T., et al., Dlx1/2 are Central and Essential Components in the Transcriptional Code for Generating Olfactory Bulb Interneurons. Cereb Cortex, 2019. 29(11): p. 4831–4849.

64. de Lombares, C., et al., Dlx5 and Dlx6 expression in GABAergic neurons controls behavior, metabolism, healthy aging and lifespan. Aging (Albany NY), 2019. 11(17): p. 6638–6656.

65. Wang, B., T. Lufkin, and J.L. Rubenstein, Dlx6 regulates molecular properties of the striatum and central nucleus of the amygdala. J Comp Neurol, 2011. 519(12): p. 2320–34.

66. Poitras, L., et al., The proneural determinant MASH1 regulates forebrain Dlx1/2expression through the I12b intergenic enhancer. Development, 2007. 134(9): p. 1755-1765.

67. Long, J.E., et al., Dlx1&2 and Mash1 transcription factors control striatal patterning and differentiation through parallel and overlapping pathways. The Journal of Comparative Neurology, 2009. 512(4): p. 556–572.

68. Fazel Darbandi, S., et al., Increased Sociability in Mice Lacking Intergenic Dlx Enhancers. Front Neurosci, 2021. 15: p. 718948.

69. Lindtner, S., et al., Genomic Resolution of DLX-Orchestrated Transcriptional Circuits Driving Development of Forebrain GABAergic Neurons. Cell Rep, 2019. 28(8): p. 2048–2063 e8.

70. Lee, H., et al., Cell Type-Specific Transcriptomics Reveals that Mutant Huntingtin Leads to Mitochondrial RNA Release and Neuronal Innate Immune Activation. Neuron, 2020. 107(5): p. 891–908 e8.

71. Taubes, A., et al., Experimental and real-world evidence supporting the computational repurposing of bumetanide for APOE4-related Alzheimer’s disease. Nat Aging, 2021. 1(10): p. 932–947.

72. Uenaka, T., et al., In silico drug screening by using genome-wide association study data repurposed dabrafenib, an anti-melanoma drug, for Parkinson’s disease. Hum Mol Genet, 2018. 27(22): p. 3974–3985.

73. Duan, Q., et al., L1000CDS(2): LINCS L1000 characteristic direction signatures search engine. NPJ Syst Biol Appl, 2016. 2: p. 16015-.

74. Cleren, C., et al., Celastrol protects against MPTP- and 3-nitropropionic acid-induced neurotoxicity. J Neurochem, 2005. 94(4): p. 995–1004.

75. Dompierre, J.P., et al., Histone deacetylase 6 inhibition compensates for the transport deficit in Huntington’s disease by increasing tubulin acetylation. J Neurosci, 2007. 27(13): p. 3571–83.

76. Hockly, E., et al., Suberoylanilide hydroxamic acid, a histone deacetylase inhibitor, ameliorates motor deficits in a mouse model of Huntington’s disease. Proc Natl Acad Sci U S A, 2003. 100(4): p. 2041–6.

77. Joshi, T., et al., Withaferin A Induces Heat Shock Response and Ameliorates Disease Progression in a Mouse Model of Huntington’s Disease. Mol Neurobiol, 2021. 58(8): p. 3992–4006.

78. Zhang, Y.Q. and K.D. Sarge, Celastrol inhibits polyglutamine aggregation and toxicity though induction of the heat shock response. J Mol Med (Berl), 2007. 85(12): p. 1421–8.

79. Lo, C.H., et al., Discovery of Small Molecule Inhibitors of Huntingtin Exon 1 Aggregation by FRET-Based High-Throughput Screening in Living Cells. ACS Chem Neurosci, 2020. 11(15): p. 2286–2295.

80. Wang, Z., et al., L1000FWD: fireworks visualization of drug-induced transcriptomic signatures. Bioinformatics, 2018. 34(12): p. 2150–2152.

81. Frey, R.R., et al., Trifluoromethyl ketones as inhibitors of histone deacetylase. Bioorg Med Chem Lett, 2002. 12(23): p. 3443–7.

82. Heyninck, K., et al., Withaferin A inhibits NF-kappaB activation by targeting cysteine 179 in IKKβ. Biochemical pharmacology., 2014. 91(4): p. 501–509.

83. Lee, H.H., et al., LPS-induced NFkappaB enhanceosome requires TonEBP/NFAT5 without DNA binding. Sci Rep, 2016. 6: p. 24921.

84. Oh, J.H., et al., Withaferin A inhibits iNOS expression and nitric oxide production by Akt inactivation and down-regulating LPS-induced activity of NF-κB in RAW 264.7 cells. European journal of pharmacology., 2008. 599(1-3): p. 11–17.

85. Oh, J.H. and T.K. Kwon, Withaferin A inhibits tumor necrosis factor-α-induced expression of cell adhesion molecules by inactivation of Akt and NF-κB in human pulmonary epithelial cells. International immunopharmacology., 2009. 9(5): p. 614–619.

86. Tanaka, K., et al., Targeting Aurora B kinase prevents and overcomes resistance to EGFR inhibitors in lung cancer by enhancing BIM- and PUMA-mediated apoptosis. Cancer Cell, 2021. 39(9): p. 1245–1261 e6.

87. Giralt, A., et al., Conditional BDNF release under pathological conditions improves Huntington’s disease pathology by delaying neuronal dysfunction. Mol Neurodegener, 2011. 6(1): p. 71.

88. Xie, Y., M.R. Hayden, and B. Xu, BDNF overexpression in the forebrain rescues Huntington’s disease phenotypes in YAC128 mice. J Neurosci, 2010. 30(44): p. 14708–18.

89. Garcia, V.J., et al., Huntington’s Disease Patient-Derived Astrocytes Display Electrophysiological Impairments and Reduced Neuronal Support. Frontiers in neuroscience., 2019. 13.

90. Mehta, S.R., et al., Human Huntington’s Disease iPSC-Derived Cortical Neurons Display Altered Transcriptomics, Morphology, and Maturation. Cell Reports, 2018. 25(4): p. 1081–1096.e6.

91. Evangelista, J.E., et al., SigCom LINCS: data and metadata search engine for a million gene expression signatures. Nucleic Acids Res, 2022. 50(W1): p. W697–709.

92. Brzezinski, J.A., et al., Ascl1 expression defines a subpopulation of lineage-restricted progenitors in the mammalian retina. Development, 2011. 138(16): p. 3519–3531.

93. Baydyuk, M. and B. Xu, BDNF signaling and survival of striatal neurons. Frontiers in cellular neuroscience., 2014. 8.

94. Battaglia, G., et al., Early defect of transforming growth factor β1 formation in Huntington’s disease. Journal of Cellular and Molecular Medicine, 2011. 15(3): p. 555–571.

95. Smith-Dijak, A.I., M.D. Sepers, and L.A. Raymond, Alterations in synaptic function and plasticity in Huntington disease. Journal of Neurochemistry, 2019. 150(4): p. 346–365.

96. Park, H. and M.-M. Poo, Neurotrophin regulation of neural circuit development and function. Nature Reviews Neuroscience, 2013. 14(1): p. 7–23.

97. Little, J.L., et al., Inhibition of Fatty Acid Synthase Induces Endoplasmic Reticulum Stress in Tumor Cells. Cancer research., 2007. 67(3): p. 1262–1269.

98. Chandwani, S., et al., Induction of DARPP-32 by Brain-Derived Neurotrophic Factor in Striatal Neurons In Vitro Is Modified by Histone Deacetylase Inhibitors and Nab2. PLoS ONE, 2013. 8(10): p. e76842.

99. Guo, Z., et al., Striatal neuronal loss correlates with clinical motor impairment in Huntington’s disease. Movement Disorders, 2012. 27(11): p. 1379–1386.

100. Nguyen, K.Q., V.V. Rymar, and A.F. Sadikot, Impaired TrkB Signaling Underlies Reduced BDNF-Mediated Trophic Support of Striatal Neurons in the R6/2 Mouse Model of Huntington’s Disease. Frontiers in cellular neuroscience., 2016. 10.

101. Bibb, J.A., et al., Severe deficiencies in dopamine signaling in presymptomatic Huntington’s disease mice. Proceedings of the National Academy of Sciences, 2000. 97(12): p. 6809–6814.

102. Bae, B.-I., et al., p53 Mediates Cellular Dysfunction and Behavioral Abnormalities in Huntington’s Disease. Neuron, 2005. 47(1): p. 29–41.

103. Mielcarek, M., et al., HDAC4 Reduction: A Novel Therapeutic Strategy to Target Cytoplasmic Huntingtin and Ameliorate Neurodegeneration. PLoS Biology, 2013. 11(11): p. e1001717.

104. Lawrence, M., et al., Software for computing and annotating genomic ranges. PLoS Comput Biol, 2013. 9(8): p. e1003118.

105. Love, M.I., W. Huber, and S. Anders, Moderated estimation of fold change and dispersion for RNA-seq data with DESeq2. Genome Biol, 2014. 15(12): p. 550.

106. Huang, R., et al., The NCATS BioPlanet - An Integrated Platform for Exploring the Universe of Cellular Signaling Pathways for Toxicology, Systems Biology, and Chemical Genomics. Front Pharmacol, 2019. 10: p. 445.

107. Chen, E.Y., et al., Enrichr: interactive and collaborative HTML5 gene list enrichment analysis tool. BMC Bioinformatics, 2013. 14(1): p. 128.

108. Korsunsky, I., et al., Fast, sensitive and accurate integration of single-cell data with Harmony. Nature Methods, 2019. 16(12): p. 1289–1296.

