## Supplementary figures and images for "Transcriptomic Characterization Reveals Disrupted Medium Spiny Neuron Trajectories in Huntington’s Disease and Possible Therapeutic Avenues"

### Supplemental Figure 1-4

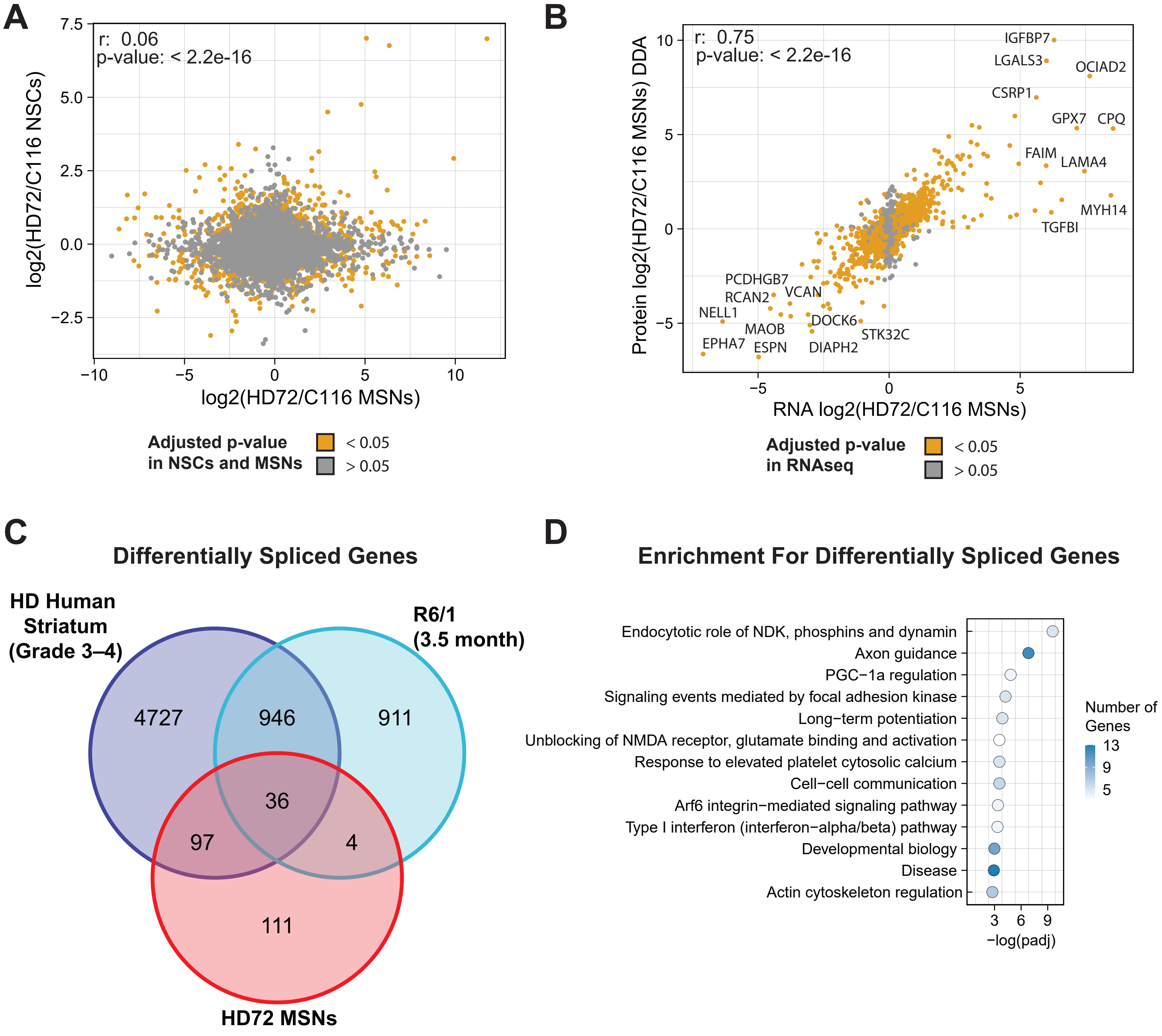


**Supplemental Figure 1**

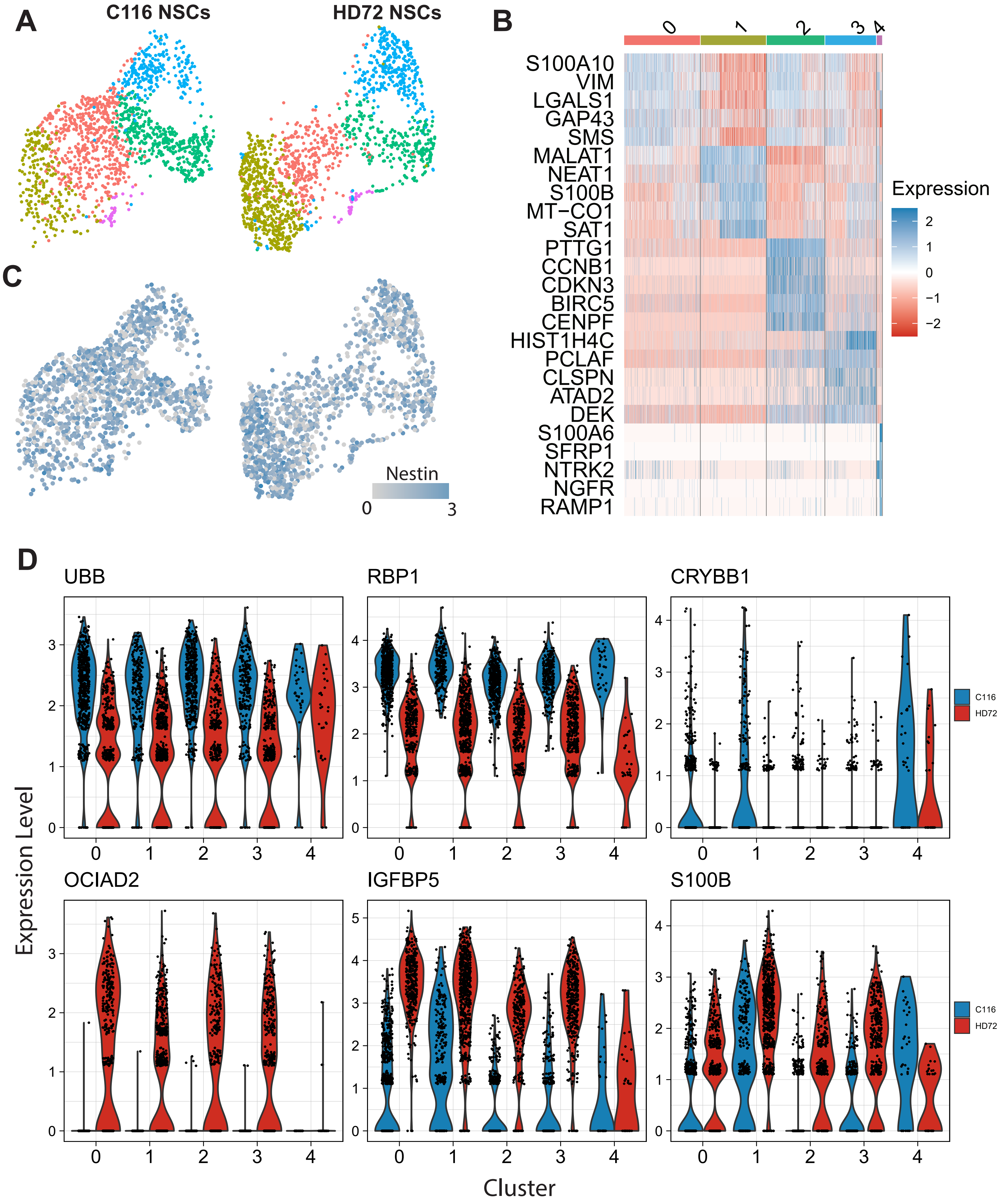


**Supplemental Figure 2**

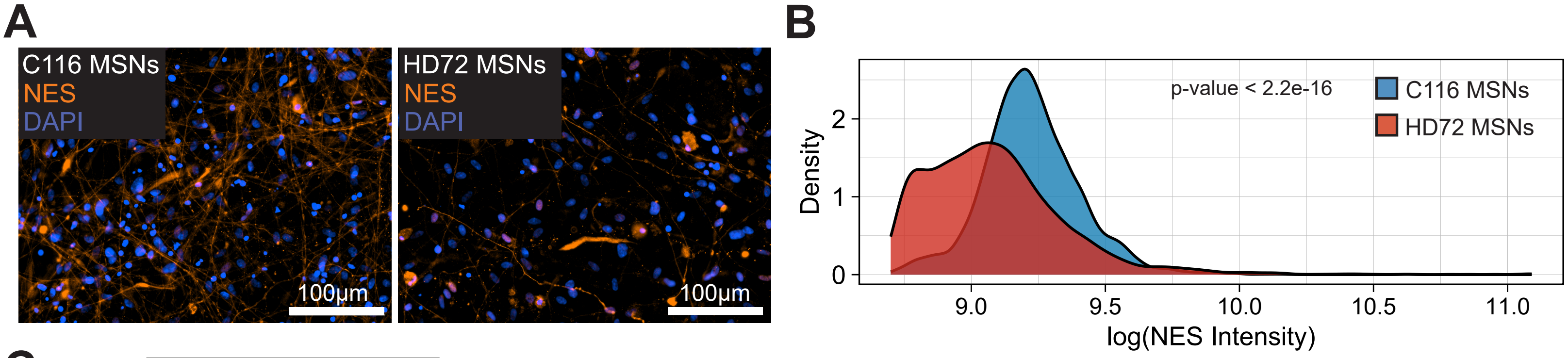


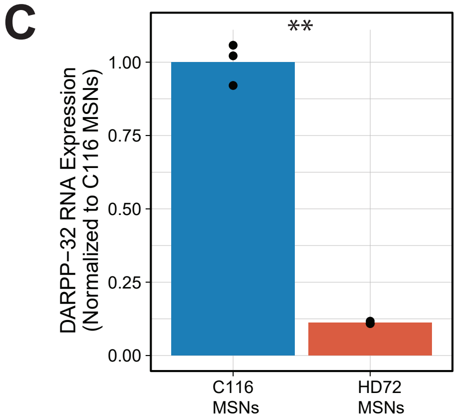


**Supplemental Figure 3**

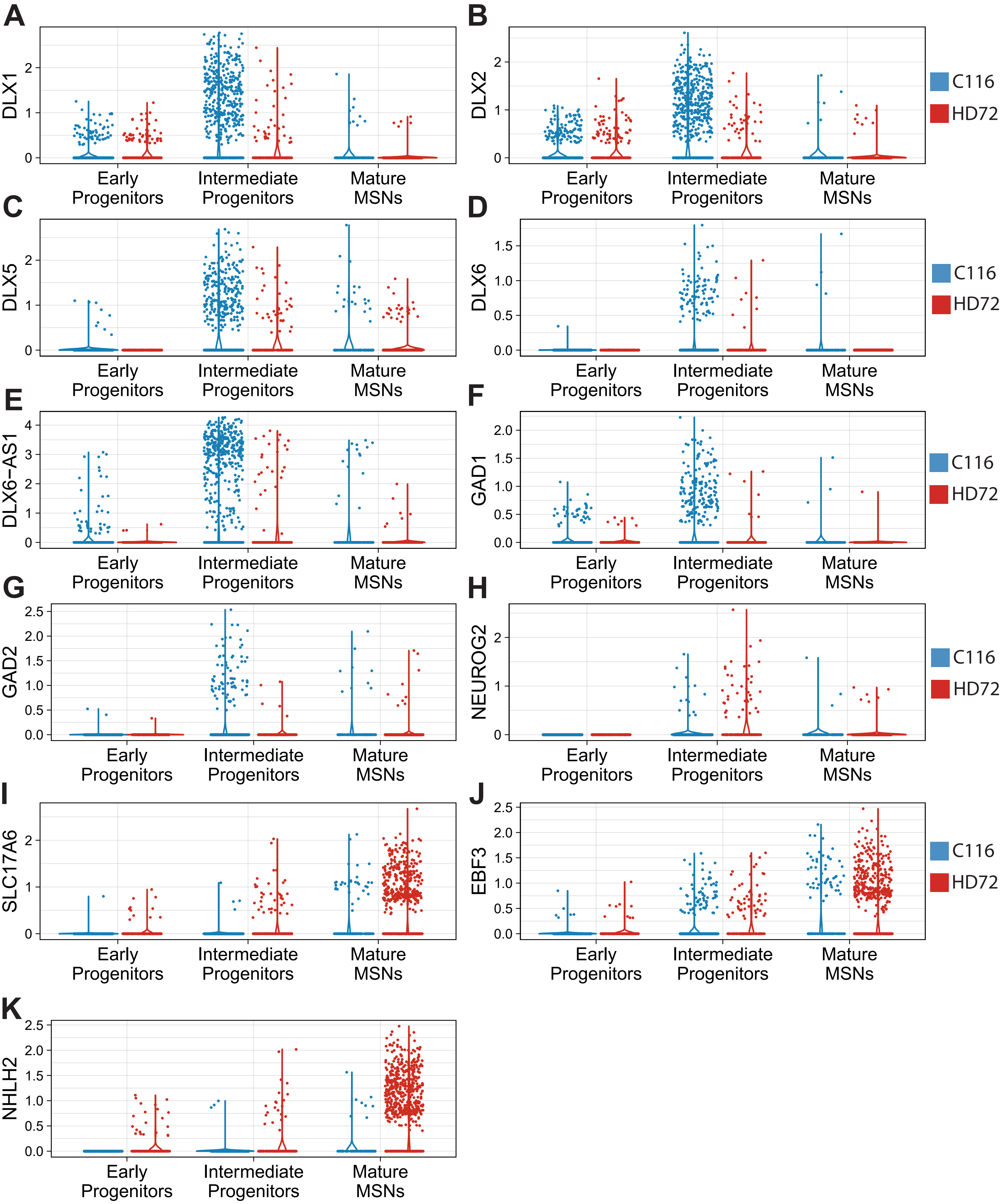


**Supplemental Figure 4**
